# A humanized mouse model system mimics prenatal Zika infection and reveals premature differentiation of neural stem cells

**DOI:** 10.1101/2025.02.21.639556

**Authors:** Allison R. Horvath, Clara M. Abdelmalek, Eunbin Park, Aubrey P. Alexander, Sadhana A. Maheswaran, Arnav H. Patel, Nandi G. Patel, Janet E. Ruan, Ademide T. Adeyemo, Erin C. Li, Katherine E. Helmicki, Stephen Lin, Paul C. Wang, Zhen Li, Li Wang, Heather A. Gordish-Dressman, Tarik F. Haydar, Tamer A. Mansour, Youssef A. Kousa

## Abstract

Zika, a mosquito-borne flavivirus, has been found in 87 countries and territories. Global outbreaks peaked in 2016. Prenatal infection of Zika virus was found to be associated with microcephaly, arthrogryposis, intracranial calcifications, fetal growth restriction, and fetal demise. The most severely affected children were diagnosed with congenital Zika syndrome, which impacts thousands worldwide. With no approved treatment or preventative measures for Zika, future viral outbreaks have the potential to cause epidemic levels of prenatal brain injury, as seen over the past 70 years. Therefore, there is a great need for a reliable and clinically translational experimental system that mimics the human condition of prenatal Zika infection. To this end, we developed a humanized, immunocompetent mouse model system of virally induced brain injury from prenatal Zika infection, which ranges from mild to severe. Here, we describe the extent to which this system mirrors the human phenotypic spectrum. Using our thorough preclinical system, we find that prenatal Zika infection of mice impacts survival rate, anthropometric measurements, tissue formation, and neurological outcomes, all of which are typical of prenatal infection. Single-cell RNA sequencing of the Zika-infected cerebral cortex reveals severely disrupted transcriptome profiles and suggests that these injuries are a result of a depletion of neural stem cells. Current and future applications include the identification of genetic or environmental modifiers of brain injury, molecular or mechanistic studies of pathogenesis, and preclinical evaluation of future therapies.

## INTRODUCTION

Prenatal viral infections can cause severe damage to the developing brain. Among the most threatening prenatal pathogens, the Zika virus is a positive-stranded RNA flavivirus, mainly transmitted by the Aedes mosquito. It was originally isolated in 1947 from a Rhesus monkey in Uganda. By 2015, it had reached Africa, Asia, and the Americas, where it caused an epidemic.^1^ Shortly after, it was shown that Zika can cross the placental and fetal blood-brain barriers during pregnancy, causing adverse outcomes for a developing fetus.^2^ This constituted a Public Health Emergency of International Concern, as declared by the World Health Organization (WHO).^3^ Severe outcomes after prenatal Zika exposure can include microcephaly, ocular abnormalities, congenital contractures, seizures, and fetal demise.^4^ While the severity of outcomes after prenatal viral infection can vary, the pattern of phenotypes in the most severely affected infants is termed congenital Zika syndrome. Currently, there is no standard of care or treatment for prenatal Zika infection, irrespective of degree of injury or severity of outcome. Further, lower levels of Zika transmission currently reduce herd immunity and increase risk for large outbreaks in the future.^1,5^ Hence, there is a pressing need for translational experimental systems of prenatal infection that effectively mimic viral propagation, brain development and injury, and host defense mechanisms.

Mouse models are integral in studying and treating viral diseases. However, the mouse immune response to Zika virus does not mimic that of humans. Unique to human infection is the ability for Zika virus to evade the interferon-I immune response. A nonstructural viral protein, NS5, binds to and degrades the STAT2 protein, a signal transducer of transcription for interferon-stimulated genes.^6,7^ However, in mice, this interaction does not occur, and the interferon-I immune response degrades the virus, preventing its propagation to the fetus. Some studies have overcome this barrier by suppressing the murine immune system, modifying interferon signaling, or inoculating virus directly into the neonatal mouse brain.^8,9^ Such studies have demonstrated several features of human infection, including maternal and fetal viremia,^10,11^ fetal resorption,^12^ fetal growth restriction,^11,13^ brain malformations,^11,13^ and ocular defects^13^. Similarly, studies in immunocompetent mice have furthered our understanding of such features, including maternal and fetal viremia,^14–16^ placental injury,^14,17^ fetal resorption,^14,15,17,18^ fetal growth restriction,^15^ brain malformations,^16,19^ neuronal loss,^16,20^ ocular defects,^16^ motor function deficits,^21^ and paralysis.^16,21^

Although each model is informative, these systems do not replicate the full spectrum of outcomes seen after prenatal viral infection or reflect the natural disease pathophysiology. In humans, these outcomes fall on a continuous spectrum, ranging from infants who are unaffected to those who appear normal at birth but later develop delays, and extending to those who suffer from severe neurological deficits. To date, a translational system that replicates the biological variability seen in humans, including fetal pathology, natural history, and developmental outcomes, has not been demonstrated.^8^ Such a system is needed to understand viral pathogenesis and evaluate candidate therapies for preventing prenatal viral infection and fetal injury.

To address these issues, we developed an experimental system that replicates viral teratogenesis after prenatal Zika infection. We used a knock-in (KI) mouse model, created by Gorman et al., 2018, in which the human STAT2 (hSTAT2) gene was inserted into the mouse *Stat2* locus via homologous recombination.^22^ The researchers demonstrated that prenatal infection of these humanized mice resulted in placental infection and fetal viremia. With this murine allele, we thoroughly evaluated the impact of infection on mice by examining the extent to which the system mimicked prenatal infection.

Here, using a novel immunocompetent humanized mouse model and an approach that mimics human exposure, we developed an experimental system that includes several characteristics pertinent to prenatal viral infections, including: 1) vertical transmission of virus from dam to fetus,^23,24^ 2) detection of virus in the developing brain,^25,26^ 3) dose- and time-dependent brain injury and fetal growth restriction,^27–29^ 4) global and regional brain injury, as evaluated prenatally and postnatally,^24,25,30,31^ 5) developmental delays in the postnatal period,^32,33^ 6) mild neurological deficits in neurological outcomes,^33,34^ and 7) overall variability in trajectory, ranging from mild neurological delays to demise.^33,35,36^ Molecular profiling of the cerebral cortex identified premature terminal differentiation of neural stem cells as the likely cause of severe brain injury and neurological deficits.

Below, we describe the system, in which hSTAT2^KI/KI^ pregnant dams were infected at timepoints developmentally analogous to the end of the first trimester in humans, which is critical for both placentation and neurogenesis. In murine development, placentation is complete by E12.5^37^ and neural differentiation begins at E10.5, as the cortical preplate forms, and continues until E17.5.^38^ Thus, we elected to evaluate the effects of infection at E9.5 and E10.5, before placentation and before (E9.5) and after (E10.5) the start of differentiation. In several domains, our system mimics the human condition and sheds light on the pathogenesis of prenatal Zika infection. Importantly, these results establish the use of this preclinical system in future research on viral pathogenesis, genetic susceptibility, and testing preventative therapies for prenatal viral infection.

## RESULTS

### An experimental system of prenatal Zika infection in a humanized immunocompetent mouse, with pipelines for both prenatal and postnatal analyses

In our experimental system, hSTAT2^KI/KI^ pregnant dams were injected intraperitoneally either with a mouse-adapted Zika virus (Zika-infected) or Dulbecco’s phosphate buffered saline (DPBS, mock-infected) on embryonic day 9.5 (E9.5) or E10.5 (**Fig. 1a**).^22^ For prenatal analyses, embryos were harvested on E12.5 or E15.5 and underwent a series of evaluations, including molecular profiling, histological analysis, and anthropometric measurements. For postnatal analyses, mice were monitored for survival and weight progression following birth, assessed for the achievement of gross motor milestones and advanced neurological outcomes, and underwent whole-brain magnetic resonance imaging (MRI).

**Fig. 1:**
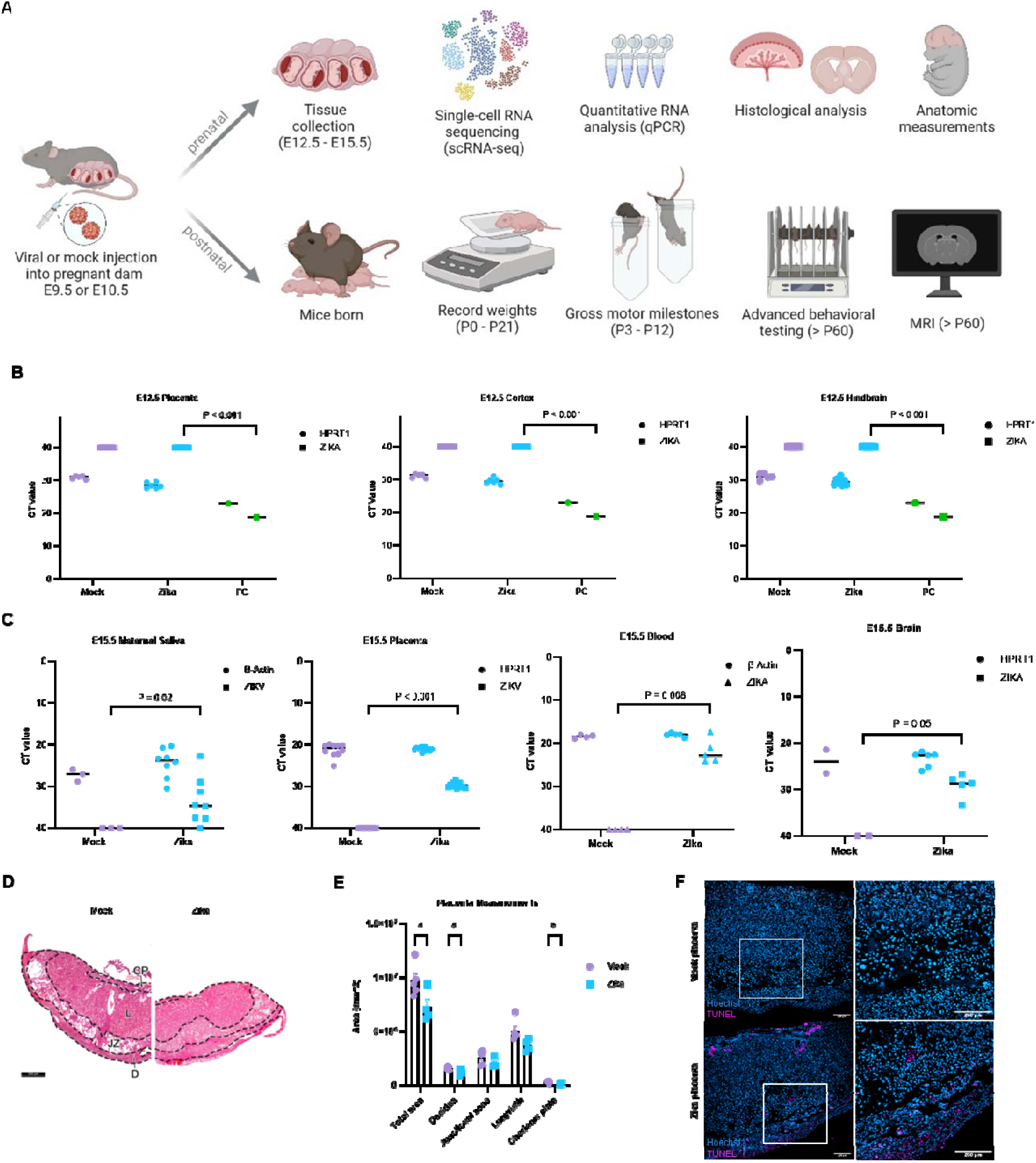
Zika evades the maternal immune system and injures the placenta. **a** Experimental design of prenatal Zika infection in a humanized immunocompetent (hSTAT2^KI/KI^) mouse model system with pipelines for both prenatal and postnatal analyses. Created in BioRender. Kousa, Y. (2025) https://BioRender.com/z27p901 **b** qPCR detects no viral RNA in the placenta, cortex, or hindbrain at E12.5 after E9.5 injection. **c** qPCR reveals viral RNA in maternal saliva, placenta, fetal blood, and developing brain at E15.5 after E9.5 infection. **d** Representative H&E image of mock- or Zika-infected placentas, with designations of regions measured in **1e**, infected at E9.5 and collected at E15.5 (scalebar is 500µm). **e** Gross measurements of placental histology confirm that infection at E9.5 causes placental injury by E15.5, most notably within the decidua and chorionic plate. **d** Immunostaining of mock- or Zika-infected placenta with TUNEL staining identifies DNA breaks resulting from apoptosis. We used (*) when P < 0.05; (**) when P < 0.01; and (***) when P < 0.001.

### Zika evades the maternal immune system and injures the placenta

#### Penetration and Injury

In humans, Zika has been found to cross the maturing placental and blood-brain barriers, localize to the fetal brain, and cause injury.^25^ As such, we performed prenatal analyses to determine viral penetration and injury in the developing brain after E9.5 infection (**Fig. 1b**). We performed qPCR on the placenta, cerebral cortex, and hindbrain for mock- or Zika-infected embryos at E12.5 and found no viral RNA compared to a positive control (PC) of pure viral cDNA. Just three days later, at E15.5, we observed amplification of Zika RNA in maternal saliva, placenta, fetal blood, and fetal whole brain, indicating maternal systemic infection and viral entry into the brain, consistent with existing literature (**Fig. 1c**).^39^

#### Placental Injury

Next, following viral pathogenesis, we evaluated the placenta of mock- or Zika-infected embryos. We found that Zika-infected embryos (E9.5) had smaller placentas, likely resulting from reductions in the decidua and chorionic plate, most notably in peripheral regions (**Fig 1d-e**). The junctional zones and labyrinth also trended smaller for Zika-infected mice, but did not achieve statistical significance. We performed TUNEL immunostaining to evaluate DNA fragmentation of apoptotic cells and confirmed the decidua and chorionic plate had more dying cells (**Fig 1f**).

### Prenatal Zika infection causes brain injury and fetal growth restriction in a dose-dependent manner

It has been shown that NS5 degrades the STAT2 protein in a dose-dependent manner, suggesting that increased viral dose causes higher rates of infection and more frequent and severe brain injury.^6^ To establish this relationship in our experimental system, we tested for an association between infectious dose and fetal tissue injury (**Fig. 2a**). We compared five doses of Zika infection to DPBS, with an infection point of E10.5 and a collection point of E15.5. We observed dose-dependent fetal growth restriction, brain volume loss, and fetal loss rates. The size of the placenta did not change with higher infectious doses. We did not find a dose-dependent response in mouse area, brain area, brain length, or hindbrain width (data not shown). Higher viral doses also led to higher prenatal lethality, observed as resorbed clumps of tissue along the uterine horns from aborted embryos (**Fig. 2b**). An intermediate infectious dose of 2.33×10^8^ PFU allowed 83% survival with concomitant brain injury and atrophy at E15.5, and was thereby chosen for additional prenatal experiments.

**Fig. 2:**
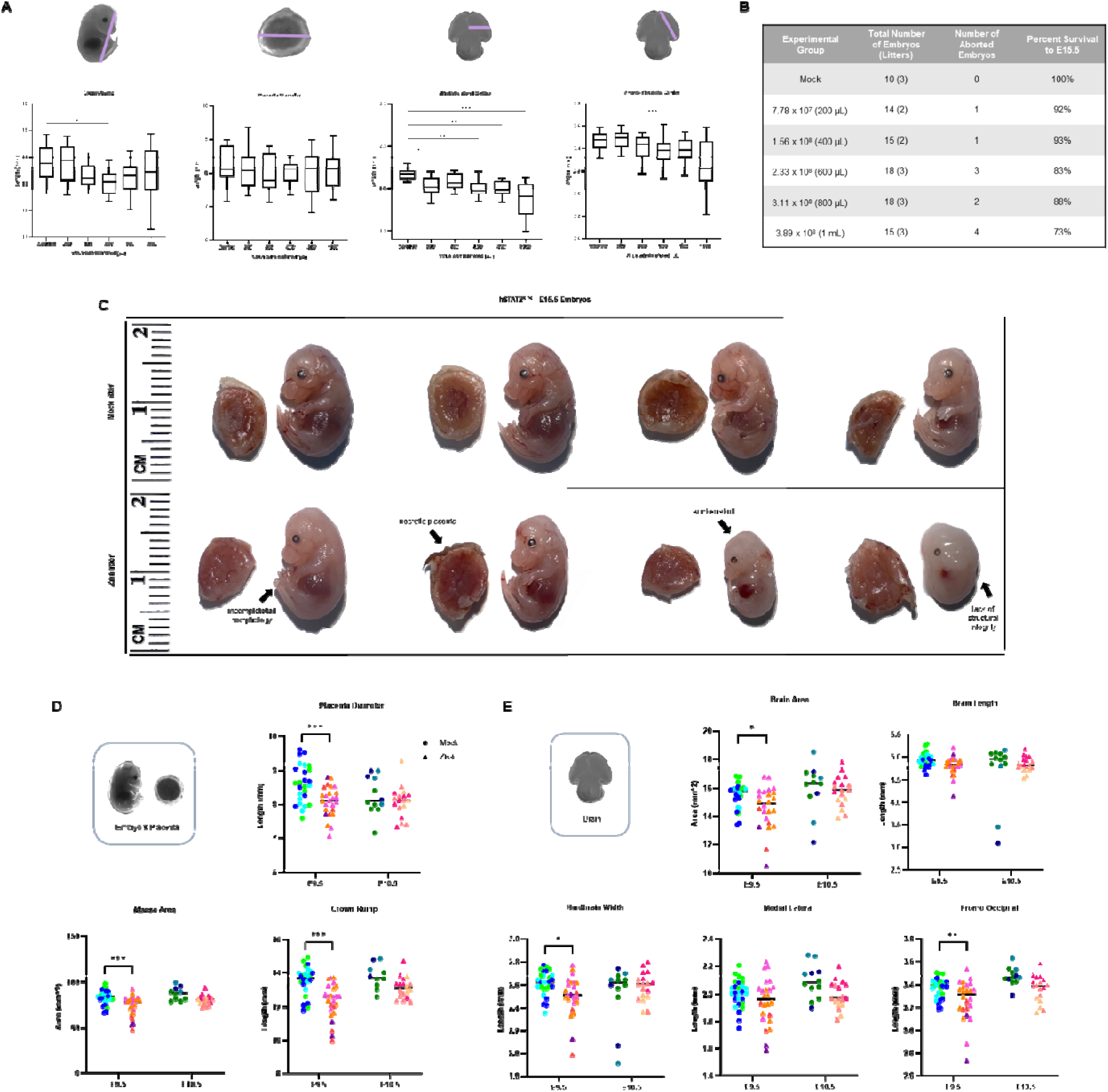
Prenatal Zika infection causes brain injury and fetal growth restriction in a dose- and time-dependent manner, on a clinically relevant spectrum. **a** A dose titration experiment, with infection at E10.5 and embryonic collection at E15.5, demonstrates a dose-dependent response of fetal growth restriction and brain volume loss. **b** Embryo loss at varying viral doses at E10.5 shows a dose-dependent response between viral dose and embryonic lethality. **c** Scaled images of mock- or Zika-infected embryos (infected at E9.5 and collected at E15.5), demonstrating variability within litters. Injuries observed in the Zika-infected litter include (from left to right embryo) an incompletely formed tail, necrosis of placenta tissue, a collapsed skull, and loss of body wall integrity. **d-e** Gross measurements of E15.5 embryos mock- or Zika-infected at E9.5 or E10.5. Each color in the figure corresponds to a different litter. There is significant fetal growth restriction, brain volume loss, and placental hypoplasia at E15.5. Earlier infection causes more severe fetal growth restriction and microcephaly. E9.5 data is based on one technical replicate. We used (*) when P < 0.05; (**) when P < 0.01; and (***) when P < 0.001.

### A single dose and timepoint of infection lead to a clinically relevant spectrum of outcomes

#### Variability in Clinical Outcomes

There are noted differences in how a prenatal viral infection can affect humans, with some infants having mild delays by five to six years of age and others having severe brain injury and neurological deficits with prenatal onset.^33,35,36^ To determine if our experimental system produced a similar spectrum of outcomes, we evaluated how a single viral dose and time of infection affected embryos and led to variability in prenatal injury. We injected pregnant dams with 2.33×10^8^ PFU of Zika or DPBS at E9.5 and observed the resulting injury at E15.5. We observed a spectrum of phenotypes ranging from mild to severe fetal teratogenesis among prenatally infected embryos, even when comparing littermates. For example, **Fig. 2c** shows a representative mock (top row) and Zika-infected litter (bottom row); while some embryos appeared unaffected by visual inspection, others lacked several anatomical structures and/or displayed embryonic malformations. Such abnormalities included incomplete formation of the tail, necrosis of the placenta, sunken skull shape, and lack of structural integrity of the body wall and limbs. The location of the embryo within the uterine horn did not seem to correlate with severity of injury (data not shown).

#### Time-Dependent Growth Restriction and Microcephaly

Beyond visual inspection, we performed detailed quantitative measurements to evaluate each mock- or Zika-infected embryo in our experiment. To perform this work, we developed a protocol to evaluate embryonic growth at E9.5 and E10.5 using a series of 2D measurements representative of standard human prenatal care. As seen in humans, prenatal infection at E9.5 resulted in embryos with significant growth restriction (smaller bodies), brain volume loss, and smaller placentas at E15.5 by two-way ANOVAs with mixed model (**Fig. 2d-e**). Notably, infection just one day later, at E10.5, did not cause any significant difference in embryo, brain, or placenta size between mock-and Zika-infected mice. A two-way ANOVA with a mixed model confirmed that viral infection at E9.5 caused significant volume loss in mouse area, crown-rump length, brain area, medial-lateral length, and fronto-occipital length compared to infection at E10.5. This is comparable to the human condition, where earlier infection has been associated with increased fetal teratogenicity and demise.^27^ We continued our studies with E9.5 infections to comprehensively evaluate the extent of brain injury and neurological disabilities, both prenatally and postnatally. Expected biological variability was observed within mock- or Zika-infected litters for infections at both timepoints. Data was evaluated by litter for non-linear trends due to maternal age and had a normal distribution (**Supplemental Fig. 4b**).

### Zika causes global and regional injury in the developing brain

#### Global and Regional Brain Injury

In addition to global brain atrophy leading to volume loss, prenatally exposed infants often also suffer from structure-specific injury and focal neurological deficits. To assess for these injuries in our mice, we collected data from three coronal regions of the forebrain and midbrain (anterior and posterior), representative of the cerebrum, with the most observable, consistent, and severe injury (**Fig. 3a-c**). Histological analyses showed injury in the developing brain after infection at E9.5, particularly manifested as cortical thinning, neuronal dropout in the pons, and reduced cellularity in multiple thalamic nuclei, including the lateral geniculate nucleus. To quantitatively determine if specific brain structures were more severely affected than others, we measured the area of the lateral ventricle, germinal zone, cortical plate, midbrain, and hemispheres (**Fig. 3d-f**). We found that infection results in significant structure- and region-specific volume loss affecting the cortical plate, germinal zone, midbrain, and overall hemisphere. Differences in the size of these structures were not uniform among the three brain regions evaluated. These data are consistent with clinical reports of varying susceptibility to injury for different brain regions and/or timepoints of infection. Some prior data has shown that the cortex, germinal zones, and thalamus may be especially injured after infection.^9,25,27,40^

**Fig. 3:**
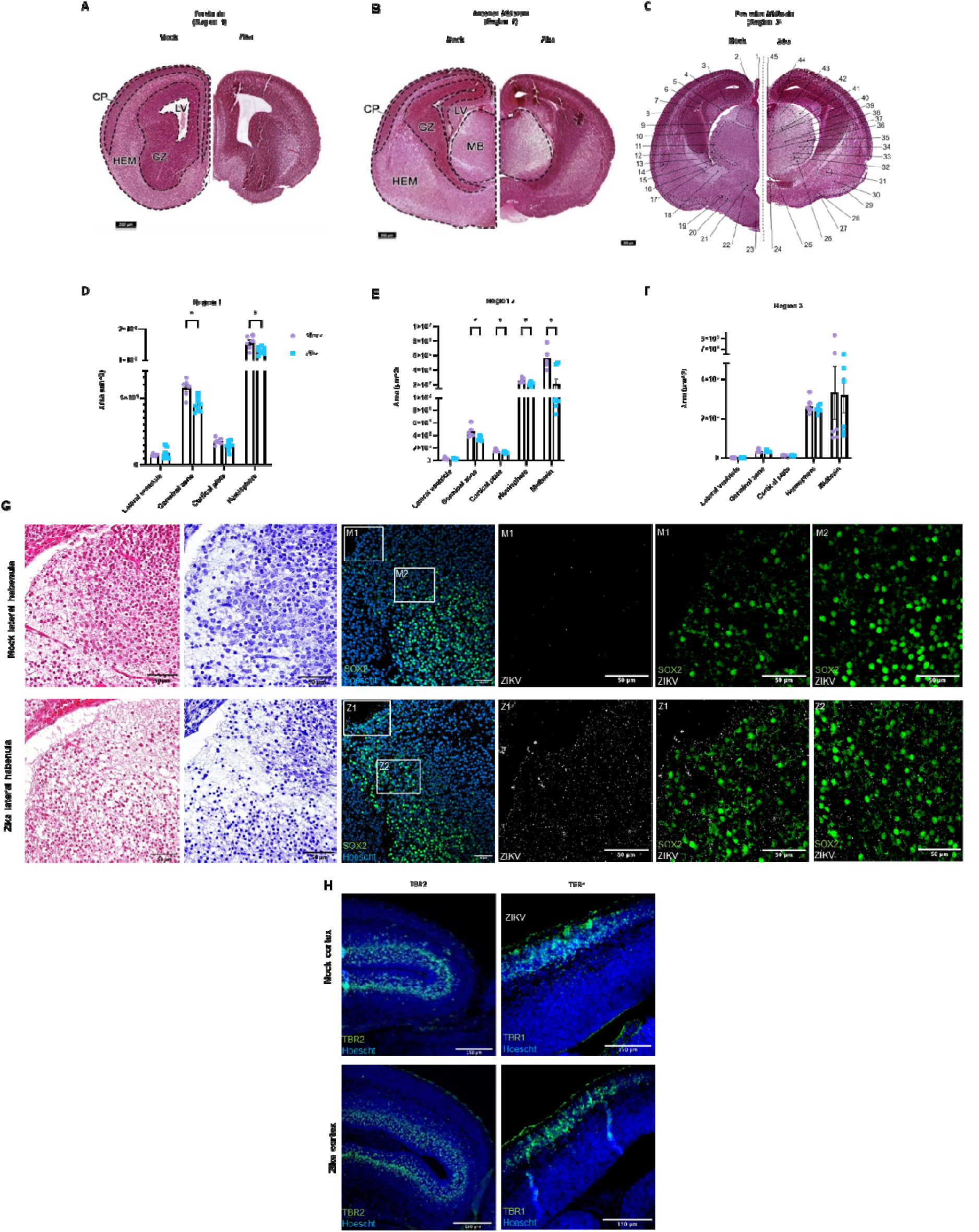
Zika causes global and regional injury in the developing brain. **a-c** Scaled images of mock- or Zika-infected brains in the forebrain (region 1), anterior midbrain (region 2), and posterior midbrain (region 3), infected at E9.5 and collected at E15.5 (scalebar is 200µm). Hematoxylin & eosin staining reveals global cerebral loss, cortical thinning, neuronal dropout in the pons, and reduced cellularity in the thalamus. **c** 1) Pineal gland, 2) Retrosplenial cortex, 3) Hippocampus, 4) Somatosensory cortex, 5) Dentate gyrus, 6) Lateral ventricle, posterior horn, 7) Choroid plexus, 8) Piriform cortex, 9) Fimbria, 10) Dorsal lateral geniculate nucleus, 11) External medullary lamina, 12) Ventral lateral geniculate nucleus, 13) Reticular thalamic nucleus, 14) Lateral amygdaloid nucleus, 15) Stria terminalis, 16) Internal capsule, 17) Lenticular fasciculus, 18) Cerebral peduncle, 19) Medial forebrain bundle, 20) Fornix, 21) Paraventricular hypothalamic nucleus, 22) Anterodorsal hypothalamic nucleus, 23) Anteroventral hypothalamic nucleus, 24) Third ventricle, 25) Reuniens thalamic nucleus, 26) Zona incerta, 27) Lateral olfactory tract, 28) Nucleus of lateral olfactory tract, 29) Cortical amygdaloid nucleus, 30) Anterior amygdaloid area, 31) Central amygdaloid nucleus, 32) Medial lemniscus, 33) Ventral medial thalamic nucleus, 34) Ventral posterior medial thalamic nucleus, 35) Ventral posterior lateral thalamic nucleus, 36) Central medial thalamic nucleus, 37) Mediodorsal thalamic nucleus, 38) Posterior thalamic nucleus, 39) Lateral posterior thalamic nucleus, 40) Paraventricular thalamic nucleus, 41) Fasciculus retroflexus, 42) Stria medullaris, 43) Lateral habenular nucleus, 44) Medial habenular nucleus, 45) Habenular commissure. **d-f** Brain area measurements of the lateral ventricle (LV), germinal zone (GZ), cortical plate (CP), midbrain (MB), and hemisphere (HEM), taken from three representative forebrain and midbrain regions. Data reveals that infection results in significant structure and region-specific volume loss affecting the cortical plate, germinal zone, midbrain, and overall hemisphere. **G** H&E and Nissl staining of coronal sections of mock- or Zika-infected fetal brains show reduced cellularity in the lateral habenula of the midbrain. Immunostaining with a flavivirus antibody, 4G2, reveals viral protein in the midbrain. Some of this viral protein appears in neural stem cells, marked by SOX2 (scalebars are 50µm). **h** Immunostaining for TBR2 reveals a restriction in neuronal intermediate progenitors. Immunostaining for late progenitors with TBR1 is inconclusive (scalebars are 150µm).

#### Cellular Injury

Similarly, in fetal brain tissues, hematoxylin and eosin (H&E) and Nissl staining revealed reduced cellularity throughout the brain, most notably affecting the lateral habenula in both afferent (stria medullaris) and efferent (fasciculus retroflexus) pathways (**Fig. 3g**). Immunostaining showed Zika virus penetrating the midbrain, with some viral protein colocalizing with neural stem cells (marked by SOX2) by E15.5, six days after infection (**Fig. 3g Z1/Z2**). Cortical immunostaining revealed a restriction of neuronal intermediate progenitors in the subventricular zone (marked by TBR2). Staining for late progenitors (marked by TBR1) was inconclusive (**Fig. 3h**).

### Prenatal Zika infection may cause developmental delays and neurological deficits

#### Developmental Delays

Infants prenatally exposed to Zika often experience developmental delays in early childhood.^32,33^ To evaluate for these outcomes in our experimental system, we developed a postnatal analysis pipeline, adapted from an existing study (**Fig 4a**).^41^ When mice were infected with the viral dose of 2.33×10^8^ PFU at E9.5, only 1/7 pregnant dams survived their embryos to term. Therefore, we used a lower viral infectious dose (7.78×10^7^ PFU) to study postnatal development. By postnatal day 12 (P12), 62% of prenatally infected mice died (**Fig. 4b**). There is a statistically significant difference in weight for mock- or Zika-infected mice, both by comparison at each timepoint and a two-way ANOVA (**Fig. 4c**). Gross motor milestone testing between P3 - P12 (during infancy) revealed developmental delays in prenatally infected mice. Testing included the righting reflex (**4d**), locomotion (**4e**), foot angle (**4f**), hindlimb suspension (**4g**), and forelimb suspension (**4h**). At all timepoints, infected mice performed statistically worse than controls in locomotor movement. At various time points, infected mice were delayed in righting reflex and hindlimb strength compared to controls. There was no significant difference in foot angle or forelimb strength tests. Data was also evaluated by litter to evaluate for non-linear trends due to maternal age and had a normal distribution (**Supplemental Fig. 5**). To evaluate whether postnatal lethality of the most severely affected mice skewed developmental testing, we also analyzed the dataset after excluding mice that died before P12. However, neurological delays and disabilities were consistent for those that survived past P12 (data not shown).

**Fig. 4:**
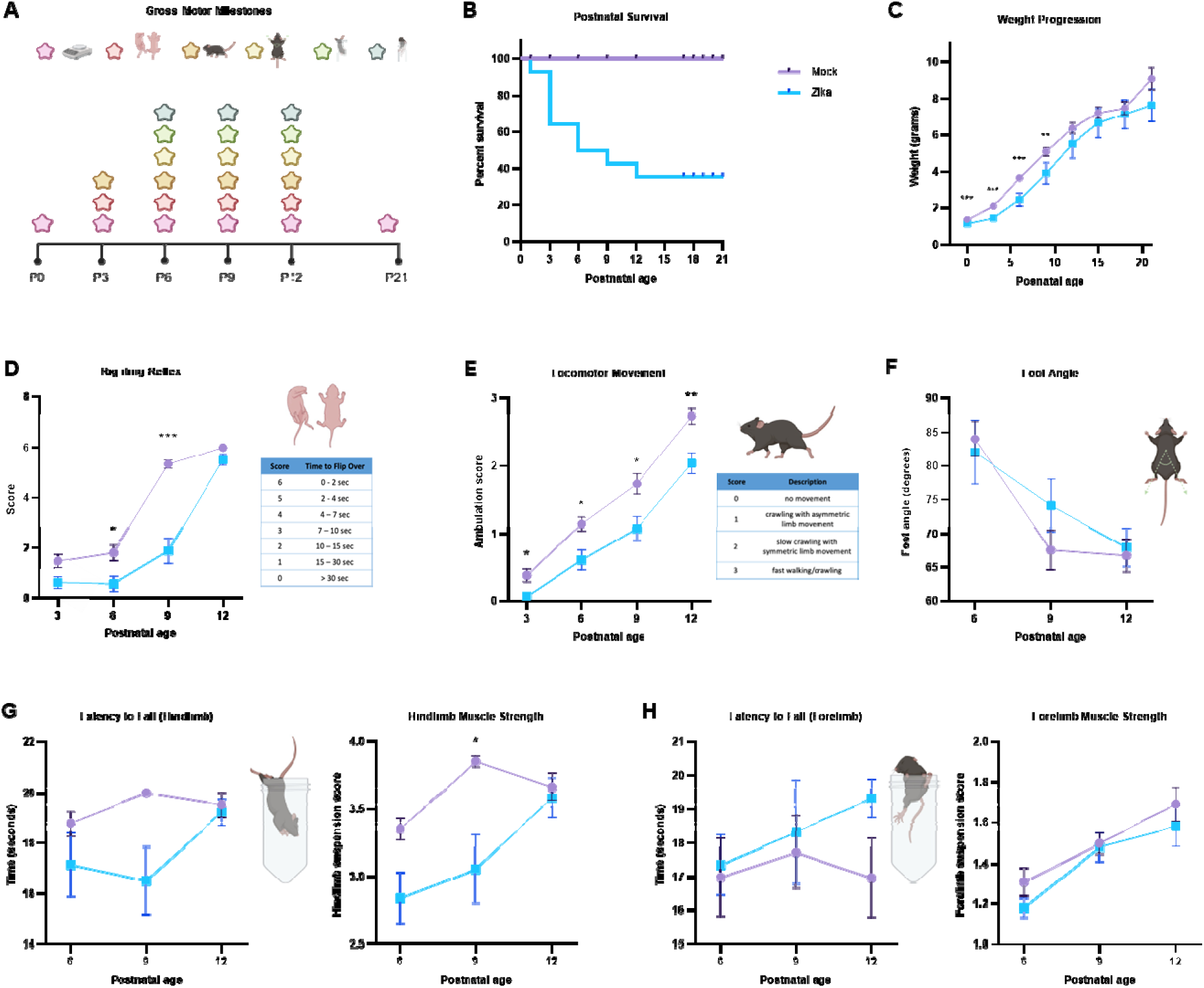
Prenatal Zika infection causes developmental delays and premature death. **a** Postnatal analysis pipeline for mice mock- or Zika-infected at E9.5. Gross motor milestone tests are performed from P3 - P12. Created in BioRender. Kousa, Y. (2025) https://BioRender.com/q46t306. **b** Kaplan-Meier curve of mock- or Zika-infected mice from P0 - P21 shows death of 62% of infected mice by P12. **c** Weight progression of mock- or Zika-infected mice from P0 - P21. Zika-infected mice consistently weighed less than controls, and there is a significant difference between the mock- or Zika-infected mice for the rate of weight gain. **d-h** Behavioral testing evaluated righting reflex (**d**), locomotor movement (**e**), foot angle while walking **(f),** hindlimb suspension **(g),** and forelimb suspension **(h).** At all timepoints, infected mice had delays in locomotor movement. At various timepoints, infected mice demonstrated delays in the righting reflex and hindlimb strength. There was no significant difference in foot angle or forelimb suspension. We used (*) when P < 0.05; (**) when P < 0.01; and (***) when P < 0.001.

#### Neurological Outcomes. Motor, Anxiety, and Social Testing

In addition to gross motor delays, prenatally infected infants also often suffer motor, social, and learning deficits later in childhood.^33,34^ We evaluated for such deficits among the 38% of mice that survived past weaning (P21) using advanced behavior tests, during both late adolescence (P35 - P60) and adulthood (> P60) (**Fig. 5a**). First, mice underwent open field testing at P35 - P40 to examine basic locomotion and anxiety-like behavior (**Supplemental Fig. 2**). No significant differences between groups were observed, which may be attributed to high variability between mice. However, some tendencies did trend differently for infected mice such as clockwise rotations, rest time, and jump count (**Fig. 5b**).

**Fig. 5:**
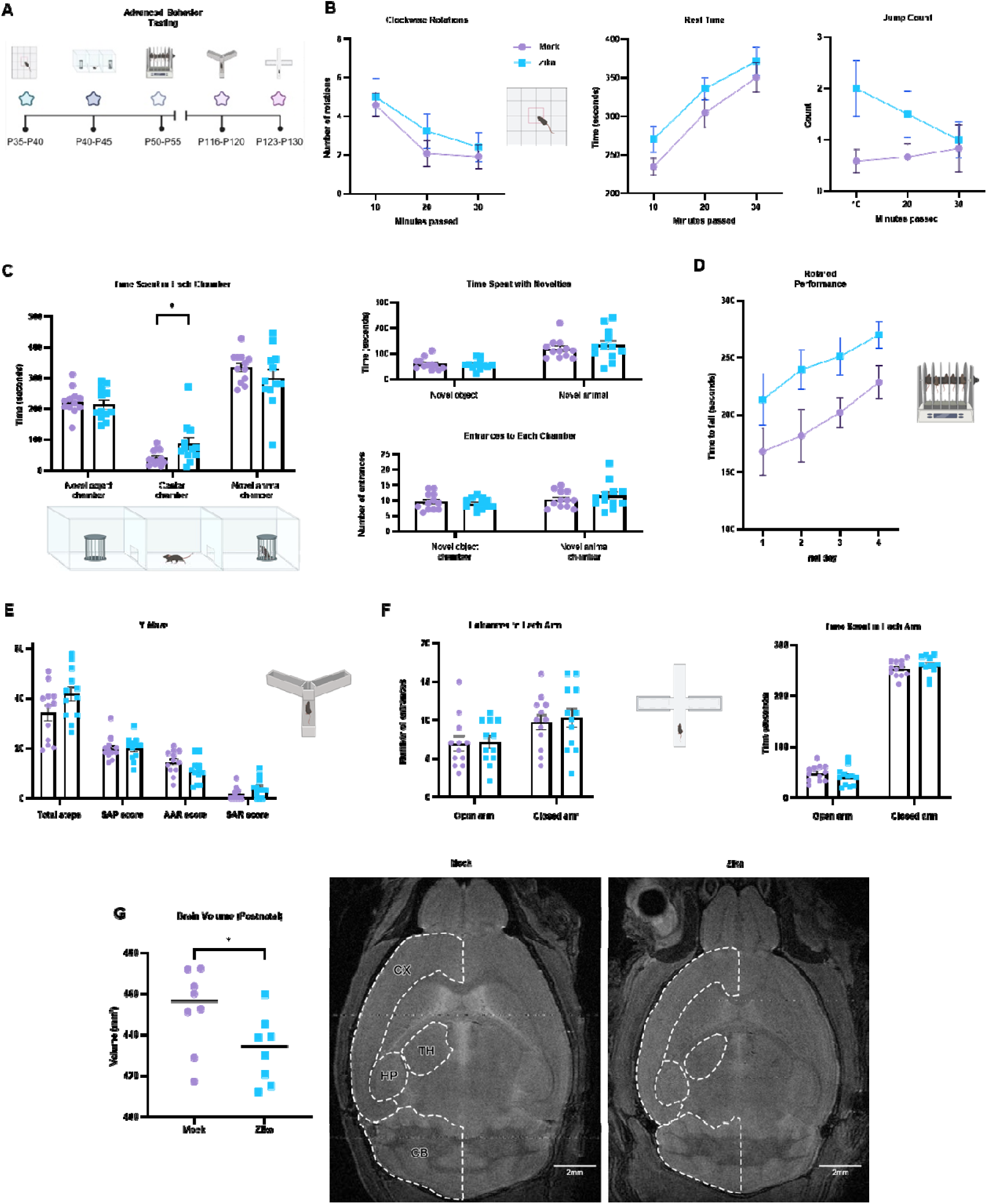
Prenatal Zika infection may cause differences in neurological outcomes. **a** Postnatal analysis pipeline for mice mock- or Zika-infected at E9.5. Advanced behavioral tests are performed from P35 - P130. Created in BioRender. Kousa, Y. (2025) https://BioRender.com/u18z535. **b** Open field test was used to examine locomotion and stereotypical movements of mice. No statistical differences were observed, although some scores, such as for clockwise rotations, rest time, and jump count trended higher for infected mice. Full dataset in **Supplemental** Fig. 2. **c** We evaluated mice for social deficits using the social interaction assay. There was no significant difference between groups in time spent with the novel object or novel animal, nor the number of entrances to either chamber. However, Zika-infected mice spent significantly more time in the empty center chamber. **d** To assess balance and motor coordination and learning, we evaluated the latency to fall from the rotarod from P50 - P55. Zika-infected mice stayed on the rotating rod longer than controls. **e** To test for differences in short-term memory, we performed the Y-Maze test. There was no significant difference in mock- or Zika-infected mice. **f** To further evaluate for anxiety, we performed the elevated plus maze test. There was no significant difference between mock- or Zika-infected mice. We used (*) when P < 0.05; (**) when P < 0.01; and (***) when P < 0.001. **g** MRI of adult brains reveals postnatal brain volume loss, potentially affecting the cortex (CX), thalamus (TH), hippocampus (HP), and cerebellum (CB) (scalebars are 2mm). We used (*) when P < 0.05; (**) when P < 0.01; and (***) when P < 0.001.

To evaluate for social deficits, mice underwent the 3-chamber socialization assay at P40 - P45, which evaluates preferences for novel objects versus novel live animals (age- and sex-matched to subject) (**Fig. 5c**). There was no significant difference observed in the time spent with each novelty or the number of entrances to each chamber. However, prenatally infected mice spent significantly more time in the center chamber (alone) than the outside chambers with the novel object or novel animal. At P50 - P55, the rotarod test was completed over four subsequent days to test both motor ability and learning potential (**Fig. 5d**). Compared to controls, prenatally infected mice remained on the rotarod for a longer amount of time before falling. There was no significant difference in the change over time in the performance of the two groups by a two-way ANOVA mixed model, indicating that there was no difference in the potential for motor learning between groups. To evaluate for short-term memory loss in prenatally infected mice, we performed the Y-Maze test from P116 - P120 (**Fig. 5e**). We observed no significant difference in short-term memory between mock- or Zika-infected mice.

Given that infants with a history of prenatal viral infection may have social-emotional deficits, it was not surprising that some of our results suggested anxious tendencies among infected mice.^42^ For example, in open field testing, some measurements trended differently between groups, such as jump count, rest time, and clockwise rotations (**Fig. 5b**). In the 3-chamber social test, greater time spent in the center chamber may have indicated anxiety in social and physical exploration. Therefore, to evaluate for anxiety, we performed the elevated plus maze test at P123 - P130 (**Fig. 5f**). However, there was no difference in time spent in enclosed (safe) versus open (exploration) arms between mock- or Zika-infected mice.

#### Postnatal Microcephaly

MRI studies of children with congenital Zika syndrome commonly report a reduction in postnatal brain volume.^4^ To characterize postnatal brain development in our model, we performed whole-brain MRI on mock- or Zika-infected mice in adulthood (**Fig. 5g**). Zika-infected mice demonstrated a significant loss in brain volume, potentially affecting the cortex (CX), thalamus (TH), hippocampus (HP), and cerebellum (CB).

### Prenatal Zika infection causes premature terminal differentiation of neural stem cells

To investigate the mechanisms underlying brain injury and adverse neurological outcomes in our model, we aimed to characterize the cellular profile of the brain prior to viral entry. To this end, we performed single-cell RNA sequencing (scRNA-seq) on seven Zika-infected cortical samples, infected at E9.5 and collected at E12.5 (**Fig. 6**). All samples were consistent in cell profile and demonstrated radial glial, neuroblast, and neuronal populations (**Fig. 6b-c**). We analyzed the distribution of cortical cells for mock- and Zika-infected samples by one-sample t-tests on parametric data and one-sample Wilcoxon sign rank tests on nonparametric data. We found that, compared to publicly available scRNA-seq datasets,^43^ which we used as our controls (E11.5, E12.5, and E13.5), our E12.5 Zika-infected sample demonstrated severe loss of 1) radial glia in the dorsal forebrain, 2) neuronal intermediate progenitors, and 3) cortical or hippocampal GABAergic neurons. In parallel, Zika-infected samples demonstrated an abundance of 1) thalamic glutamatergic neuroblasts, 2) mixed region neuroblasts (glutamatergic and general), 3) forebrain GABAergic neurons, and 4) erythrocytes (**Fig. 6d-e**). The loss of immature cells and abundance of mature neurons may suggest that Zika causes premature differentiation of neural populations in the developing brain, as seen in previous literature (**Fig. 6a**).^19,44^ Consistent with these findings, *Sox2*, an indicator of neural stem cells, was found abundant in mock-infected cells and severely restricted in Zika-infected cells (**Fig. 6f**). *Eomes*, a marker for neuronal intermediate progenitor cells, was slightly restricted in Zika-infected cells (**Fig. 6g**). *Tbr1*, largely found in mature neurons, was abundant in Zika-infected cells compared to mock-infected cells (**Fig. 6h**).

**Fig. 6:**
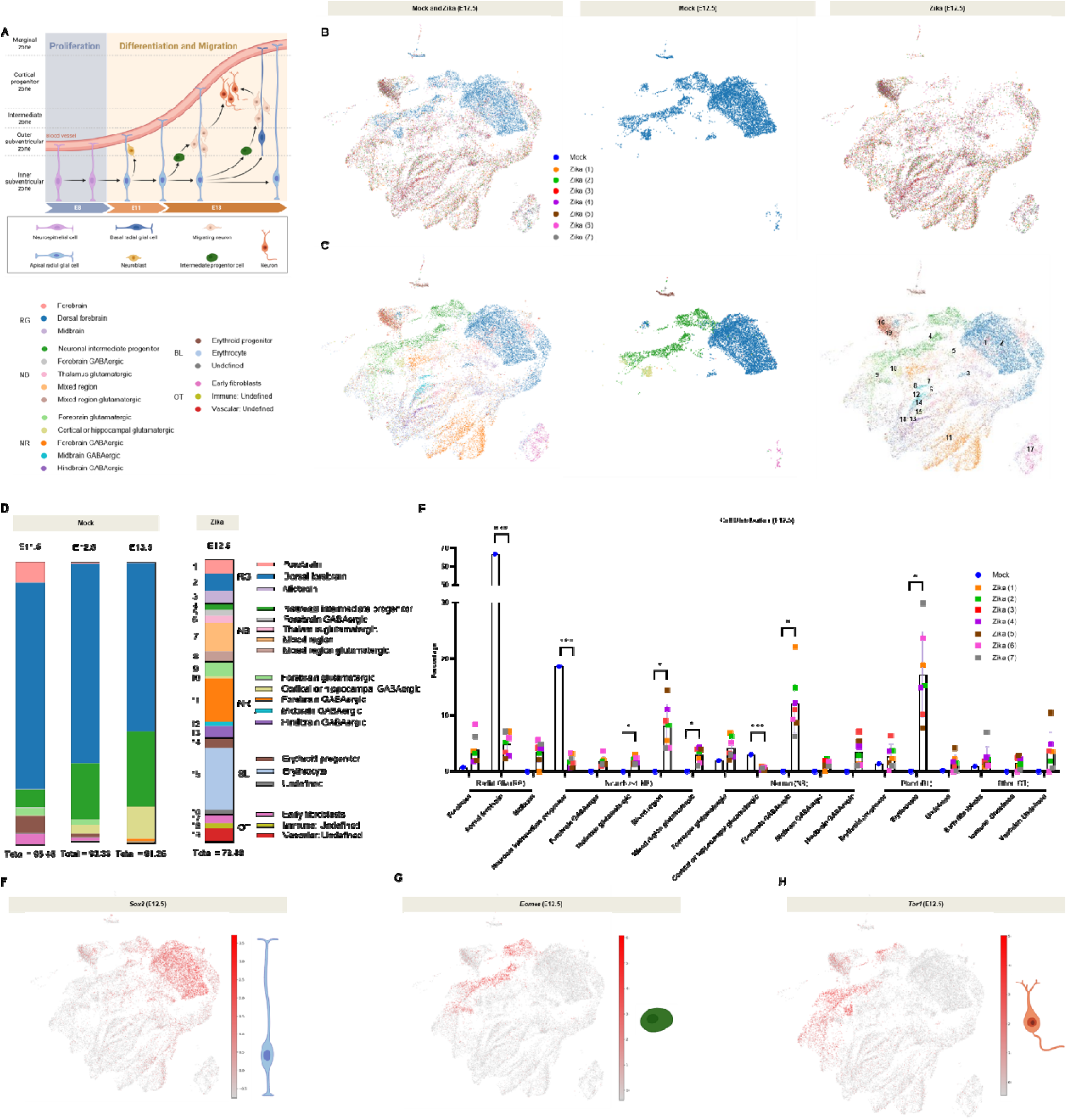
Prenatal Zika infection causes premature terminal differentiation of neural stem cells. **e** Schematic showing the development of the cortex, from neuroepithelial cells to mature neurons. Created in BioRender. Kousa, Y. (2025) https://BioRender.com/l48q033 **b-c** t-SNEs of 25,492 cortical cells, mock- or Zika-infected at E12.5, color-coded by (**b**) condition and (**c**) cell type. Radial Glia (RG): Forebrain, Dorsal forebrain, Midbrain. Neuroblast (NB): Neuronal intermediate progenitor, Forebrain GABAergic, Thalamus glutamatergic, Mixed region, Mixed region glutamatergic. Neuron (NR): Forebrain glutamatergic, Cortical or hippocampal GABAergic, Forebrain GABAergic, Midbrain GABAergic, Hindbrain GABAergic. Blood (BL): Erythroid progenitor, Erythrocyte, Undefined. Other (OT): Early fibroblasts, Immune: Undefine, Vascular: Undefined. **d-e** Distributions of mock- (E11.5, E12.5, E13.5) and Zika-infected (E12.5) cortical cells suggest that Zika causes premature terminal differentiation of neural stem cells. **f** *Sox2*, an indicator of neural stem cells, was found abundant in mock-infected cells and severely restricted in Zika-infected cells. **g** *Eomes*, a marker for neuronal intermediate progenitor cells, was slightly restricted in Zika-infected cells. **h** *Tbr1*, largely found in mature neurons, was abundant in Zika-infected cells compared to mock-infected cells. We used (*) when P < 0.05; (**) when P < 0.01; and (***) when P < 0.001.

### Prenatally infected females have enhanced postnatal survival

Outcomes of prenatal viral infection may differ by sex.^45^ We evaluated for sex-specific differences in fetal growth restriction and brain injury for the embryos infected at E9.5 and collected at E15.5 shown in **Fig. 2d-e** (**Supplemental Fig. 3**). Differences in measurements correlated to infection status, as previously shown, irrespective of sex. Further, no statistical difference in the outcomes of infection were observed between sexes by a two-way ANOVA (**Supplemental Fig. 3a-b**). However, from E15.5 to P21, the percentage of males decreased from 65% to 16%. Fisher’s Exact Test shows a statistically significant difference in distribution between sex of mock- or Zika-infected mice by P21, indicating a potential postnatal survival advantage for the female sex (**Supplemental Fig. 3c**).

#### Advanced Maternal Age

It is known that the immune system can become weaker and more susceptible to viruses with age, as with West Nile virus.^46^ In our model, we evaluate the dose titration at E10.5 shown in **Fig. 2a** by litter to determine the effects of differing maternal age (**Supplemental Fig. 4a**). Advanced maternal age contributed to adverse prenatal outcomes, particularly at higher viral doses.

## DISCUSSION

Here, we developed an immunocompetent mouse experimental system that allowed for thorough prenatal and postnatal analyses of infection, closely mirroring human outcomes. Observed phenotypes in the system encompassed both known and novel characteristics, including brain injury in the absence of virus, viral localization in the fetal brain, blood, and placenta, spontaneous abortion, fetal growth restriction, microcephaly, developmental delay, and variability in outcomes influenced by factors such as genetic differences, sex, maternal age, viral dose, and timing of infection.

In humans, Zika has been shown to cross the placental and blood-brain barriers and localize in the fetal placenta and brain.^25,39^ Three days post-infection, we did not detect virus in the placenta, cortex, or hindbrain, despite differences in neural cell development as detected by single-cell RNA sequencing. Notably, these data suggest that Zika can indirectly cause injury to the prenatal brain, potentially by weakening the maternal immune system and increasing inflammatory responses. Future work may include an investigation of indirect pathogenesis, as it may open avenues for treatment. Six days post-infection, we detected viral RNA and protein in Zika-infected embryos using qPCR and immunostaining. Placental analyses revealed virus, injury, and cell death in the decidua and chorionic plate. Notably, the decidua plays an important role in host defense mechanisms against foreign pathogens and associated congenital infections.^47^ Disruptions to the placenta may contribute to brain injury, as it contains key factors that promote neural communication and differentiation, such as brain-derived neurotrophic factor (BDNF).^48^

Although there is minimal clinical data that an increased viral dose leads to more severe outcomes in humans, it has been shown that the STAT2 protein degrades NS5 in a dose-dependent manner. Our evaluation of viral doses at E10.5 in mice revealed a dose-dependent relationship between viral exposure and fetal teratogenicity, as evidenced by demise, growth restriction, and brain atrophy and volume loss at E15.5. These data may suggest a similar relationship in humans. These experiments also identified the optimal viral dose to replicate prenatal injury while limiting the fatality of the animals. While it is not known what the average infectious dose is for humans, this work may help us identify the critical dose at which injury is sustained.

Pertinent to our design and approach is the time point of infection, which appears to correlate with injury. In humans, injury appears to be most frequent and severe for infections in the late first or early second trimester, before placentation and differentiation.^4,27,37,38^ It has been shown that the Zika virus preferentially infects undifferentiated neural stem cells (NSCs) during neurogenesis, which appear to be highly vulnerable in the developing brain.^9^ In mice, NSCs start proliferating at E8. Soon thereafter, NSCs lose certain distinguishing features, including tight junctions, and begin acquiring markers of differentiation, which can be detected by E10.5.^49^ This makes E9.5 a crucial transitional time point between neural stem cell proliferation and differentiation in some brain regions. One such region is the ventricular zone, which is comprised of neuroepithelial cells that give rise to important neural cell populations, including neurons, astrocytes, oligodendrocytes, and the ependymal cells lining the ventricular system. Therefore, we evaluated how infection during and after this transition (E9.5 vs. E10.5) would impact neuronal development. We observed that viral infection at E9.5 caused significant fetal growth restriction and microcephaly, while infection at E10.5 did not. These findings are consistent with the human condition, where earlier infection has been associated with more severe fetal teratogenicity.^27^

Various measurements of embryo, placenta, and brain demonstrated significant growth restriction or volume loss in infected mice, consistent with human outcomes.^27–29^ Notably, in humans, placental hypoplasia often contributes to fetal growth restriction.^50^ Within the Zika-infected litters, even with a single dose and timepoint of infection, we observed a spectrum of phenotypic severity, as seen in prenatal viral infections.^29,35^ These findings suggest that at least some of the differences observed among prenatally infected infants are not related to host/viral genetic or environmental differences.

Histological staining revealed reduced cellularity in the lateral habenular nucleus and thalamus, including the lateral geniculate nucleus. Importantly, the lateral habenula is involved in mood regulation and sleep, both of which may be disrupted in infants with congenital Zika syndrome.^51^ The thalamus, the sensory relay center of the brain, is involved in the regulation of cognition, vision, motion, language, and circadian rhythms, all of which are disrupted by infection.^26,52–54^ Of note, the lateral geniculate nucleus plays a key role in vision, potentially advancing our understanding of cortical visual impairment after prenatal Zika infection, as well as in other prenatal infections.^55^ These results identify specific brain structures as potential targets for future studies of therapeutics.

Immunostaining supported the notion that prenatal Zika infection causes global microcephaly by targeting neural stem cells.^9,56^ In the prenatal brain, neural stem cells are concentrated in the ventricular zone and subventricular zone. After proliferation and migration to the cortical plate, the reservoir of neural stem cells also migrates outward and localizes in the subventricular zone and dentate gyrus subgranular zone of the hippocampus.^57,58^ These areas of proliferation are known as germinal zones, both prenatally and postnatally. As Zika preferentially infects neural stem cells, it follows that the germinal zone and cortical plate were smaller after infection in our experimental system. Importantly, disruption of proliferation and cortical expansion can hinder decision-making, emotion, cognition, social interactions, and language, as seen in children with congenital Zika syndrome.^30,59–61^ A notable difference between the human condition and this system is ventriculomegaly, which is a common feature after severe infection, but not a feature of this experimental system.^40^ In humans, ventriculomegaly results from a relative enlargement of the cerebral spinal fluid spaces from brain atrophy or volume loss. In that regard, it is interesting that the experimental system demonstrated atrophy and volume loss, but not ventriculomegaly. This could be due to tissue fixation and processing, as technical variations might have influenced the size of the ventricles. On the other hand, ventriculomegaly may be a human-specific consequence, resulting from infection and inflammation of the choroid plexus. As choroid plexus is much larger in humans than it is in mice, damage to the choroid plexus in humans may cause ventricular blockage, CSF circulation defects, and ultimately ventriculomegaly.

Infants prenatally infected with Zika can also experience developmental delay and/or neurological deficits as they grow, and these deficits can shorten the lifespan of the most severely affected infants.^32–34^ In our experimental system, we observed significant delays in motor skill attainment for infected mice in the first two weeks of life. In advanced behavioral testing, infected mice showed some social differences and anxiety-like behavior, although results were not always consistent. These findings may correlate with emotional dysregulation seen in patients following prenatal Zika virus exposure.^42,62^ Due to sex-related differential survival, a large proportion of the Zika-infected mice that underwent advanced behavioral testing were female, which may have skewed results. When evaluated postnatally, 62% of Zika-infected mice died by P12.

In humans, there may be sex-specific protective or risk factors for outcomes of infection.^45^ In this experimental system, the number of males and females were nearly equal in the prenatal period after infection. In contrast, postnatally, we observed many of the males die and far more females survive to weaning age (P21). These findings suggest a sex-specific susceptibility in males or protection from infection or injury in females. The cause for this difference might be a topic for validation in human studies and a subject for future investigation in this preclinical system.

We found that advanced maternal age may be a risk factor for fetal growth restriction and microcephaly, particularly at higher viral doses. Although there is little clinical data to support this for prenatal Zika infection, it is known that advanced maternal and paternal age can contribute to adverse pregnancy outcomes.^63^

Importantly, a scRNA-seq experiment to investigate the mechanism of injury suggested that Zika causes premature terminal differentiation of neural stem cells in the developing brain, both by the depletion of neural progenitors and the abundance of inhibitory neurons. This was supported by histological data, which revealed a restriction of neuronal intermediate progenitors. Additionally, an abundance of red blood cells and a low total cell count may suggest increased cell death. Viral infection may restrict the available reservoir of proliferative cells, ultimately contributing to differential migration, microcephaly, and adverse neurodevelopmental outcomes. Future work will likely include a single-cell analysis on genetic- and age-matched controls. It may also include histological analyses to evaluate for the premature presence of inhibitory neurons and erythrocytes in the dorsal forebrain at E12.5.

Our novel, humanized, and immunocompetent experimental system replicates several characteristics pertinent to human infection, from maternal exposure and transmission through the placenta to infection of the developing brain and injury leading to a spectrum of long-term neurological deficits. As such, this system is clinically relevant and has compelling uses for studies of molecular or mechanistic studies of pathogenesis, genetic susceptibility, and in testing targeted therapies in the future.

## METHODS

### hSTAT2^KI/KI^ Colony Generation

C57BL/6-*Stat2*^tm1.1^(STAT2)^Diam^/AgsaJ mice were obtained from Jackson Labs, and genotyping was completed according to their protocol with primers AGG AGG CTG AGG TAG AAT CAC T (forward) and TGG AGA GAT GGC TCA GAG GT (reverse).^22^

### Zika Virus Strain

A mouse-adapted strain of Zika virus was used for all experiments in this study. The strain was generated by the authors of Gorman et al. by passaging an African Zika strain (ZIKV-Dak-41525) through Rag1^-/-^ mice to obtain a more virulent, mouse-adapted virus (ZIKV-Dak-MA).^22^

### Zika Virus Preparation

Vero cells were obtained from American Type Culture Collection (ATCC). At 95% confluency, all media was removed and replaced with Zika virus inoculum (5mL, MOI 0.001, 225mL tissue culture flask). The flask was continuously rocked for 10 mins, and an additional 5mL of FBS-free media was added. Next, the flask was returned to the incubator. After one hour, 45mL of media containing 2.5% FBS was added to the flask. After an additional 48 hrs, 6mL of 10X Sucrose-Phosphate-Glutamate was mixed with the cell supernatant and filtered through a 0.1μm polyethersulfone vacuum filter (Nalgene). The virus was aliquoted into 2mL cryovials, flash frozen in a dry ice/ethanol bath, and stored at -80°C. The concentration of virus was quantified by plaque forming assay.

### Zika Virus Infections

hSTAT2^KI/KI^ mice were mated, and vaginal plug was confirmed by E0.5. At E9.5 or E10.5, Zika virus or DPBS control was administered via intraperitoneal injection. For Zika infection, virus was thawed quickly at 37°C. All mice were housed in a BSL-2+ animal facility. Dams were monitored for adverse reactions after infection and throughout pregnancy. Embryos were either extracted at E12.5 or E15.5 for prenatal analyses or allowed to deliver for postnatal analyses. There was no evidence of viremia (virus in saliva and blood) at P35 and mice were moved from BSL-2+ containment to the standard housing facility for behavioral/outcome testing.

### Gross Measurements/Scoring

CellSens software was used to take images of embryos. ImageJ was used to quantify embryo, placenta, and brain measurements. All measurements were completed by two blind reviewers and averaged, with the exception of MRI data. If the difference between the two measurements differed by 25% or greater, they were analyzed by a third reviewer and averaged. Representative data is shown throughout the manuscript.

### RNA Isolation

Tissue was collected and stored in 500µL of TRIzol. Next, 100µL of phenol-chloroform was added, and tissues were disrupted using a vortex. Samples were centrifuged for 15 mins at 12,000g. Next, 100µL of the aqueous layer was removed and transferred to a round bottom Qiagen tube with 250µL lysis solution. Samples were run on the QIAcube using the *RNeasy* protocol.

### cDNA/qPCR

cDNA was prepared using the High-Capacity cDNA Reverse Transcription Kit from ThermoFisher with RNA diluted to 12.5ng/µL (E12.5) or 40ng/µL (E15.5). Some Zika quantification was completed with TaqMan Fast Advanced Master Mix using Probes from Gorman et al. and β-actin as a reference gene.^22^ Some Zika quantification was performed with Applied Biosystems Fast SYBR Green Master Mix, using HPRT1 as a reference gene. E12.5 samples were amplified an extra 25 times before qPCR to improve signal detection. All qPCR was completed on a QuantStudio 7 Pro Real-Time PCR System.

### scRNA-seq Sample Preparation

The cerebral cortex was collected from seven Zika-infected samples. Tissue was dissociated using the Neural Tissue Dissociation Kit (Miltenyi Biotec), with modifications to the standard protocol. HBSS was replaced with a custom FANS solution (0.5g BSA powder, 100µL 0.5M EDTA, and PBS) to reduce cell adhesion, and the centrifugation step was set to 300g for 5 minutes. After collection, single-cell RNA sequencing was performed using the Chromium Next GEM Single Cell 5’ Reagent Kit v2 (10x Genomics).

### scRNA-seq Data Analysis

For each sample, 10,000 cells were targeted for sequencing to achieve 50,000 paired-end fragment depth per cell. Libraries were sequenced according to the manufacturer’s instructions on the Illumnia NovaSeq X Plus 25B flow cell.

The scRNA-seq data were pre-processed using 10x Genomics CellRanger v8.0.1.^64^ Initially, the sequencing data were aligned to the mouse GRCm39 (GENCODE vM33) reference genome, supplemented by the ZIKV Dakar (Senegal, 1984, GenBank: MG758785.1) genome. Then, cellular barcodes were identified, and we performed UMI counting. The raw feature-barcode matrices were processed with the Cellbender v0.3.2 software package to remove ambient RNA molecules and correct counts affected by random barcode swapping.^65^

Next, barcode rank plots were generated using the Scanpy^66^ and Plotly^67^ packages. The UMI count at the first cliff in each sample was manually identified and used as a cutoff to filter out cells will low UMI counts. To ensure data quality, cells were excluded if they met any of the following criteria: (1) greater than 25% mitochondrial content, or (2) outliers in the population, deviating by more than five-fold from the mean absolute deviation in any of these metrics: total UMI count, UMI count of the top 20% genes, or total gene count. Additionally, cell duplicates were identified and removed by the BoostClassifier function from the doubletdetection python package.^68^

Single-cell RNA sequencing samples from a developing mouse neocortex at E11.5, E12.5, and E13.5 were obtained from the GEO SuperSeries GSE1531646 to serve as controls.^43^ The cells were subjected to the same quality filters applied to the Zika-infected cells.

All samples, including the three control samples and seven Zika-infected samples, were merged by outer joining of gene IDs. Any genes not expressed in at least 10 cells were excluded. To correct for variability in sequencing depth, the counts in all cells were normalized by library size and log2 transformed. The celltypist python package was used to annotate the tested cell types using the built-in “Developing Mouse Brain” model.^69^ The cell annotations in absolute count and percentage per sample were exported in Excel sheets.

For visualization on UMAPs, only the E12.5 control sample is included for direct comparison with the E12.5 Zika-infected samples. We used Scanpy’s ’seurat_v3’ flavor algorithm, a robust method for selecting genes informative for distinguishing cell types, to identify the 2,000 most pertinent genes. These included key markers of brain development, like *Tbr1*, *Nes*, and *Eomes*. Additional key markers, like *Sox1* and *Sox2*, were manually added. Subsequently, the expression data was zero-centered and scaled to unit variance using Scanpy. We use the harmonypy python package to remove batch effects between samples while preserving biological variation.^70^ Principal component analysis (PCA) was applied, and UMAP visualization was used to project the high-dimensional data into two-dimensional space, providing insights into the relationships between clusters. The cell populations with more than 1% of the total cell counts were kept for visualization.

### Tissue Preparation

Brain and placenta tissue were harvested and placed in 4% paraformaldehyde for 24 hrs prior to placement into PBS. Tissue was embedded by the Children’s National Hospital Pathology Core. Tissue was sectioned with a microtome into 7µm slices for histological analyses.

### H&E Staining

Sectioned tissue was deparaffinized, hydrated, stained with hematoxylin and eosin for nuclei and cytoplasm visualization, dehydrated, and cleared. Slides were mounted with an organic mounting media and #1 cover slips. A Leica DM6B microscope was used to image samples at 10x, as well as 40x for areas of interest.

### Nissl Staining

Sectioned tissue was deparaffinized, hydrated, stained with cresyl violet solution for nuclei visualization, dehydrated, and cleared. Slides were mounted with an organic mounting media and #1 cover slips. A Leica DM6B microscope was used to image samples at 10x, as well as 40x for areas of interest.

### Immunohistochemistry

Tissue was deparaffinized with xylene and rehydrated. Antigen retrieval was completed using Sodium Citrate Buffer (10mM Sodium Citrate, with or without 0.05% Tween 20, pH 6.0). Blocking was completed using goat serum or bovine serum albumin (BSA), and antibodies were diluted in blocking solution. Nuclei were stained with Hoescht 33342. Virus was stained with Monoclonal Anti-Flavivirus Group Antigen, Clone D1-4G2-4-15 (produced in vitro, VR-1852). The Click-iT™ Plus TUNEL Assay Kit was used for in situ apoptosis detection. TBR2 was stained by Anti-TBR2/Eomes antibody [EPR19012]. TBR1 was stained by Anti-TBR1 antibody [EPR8138(2)].

### P3 - P12 Gross Motor Milestone Testing

Animals were videotaped performing set tasks between P3 - P12. Videos were renamed to deidentify condition and distributed to lab members for blinded evaluation and scoring. Scores from each test were averaged from two trained scorers, unless their scores deviated more than one point and/or 2 seconds, at which point lab members re-scored that video for validation prior to final averaging.

### Righting Reflex

The righting reflex is a component of sensory-motor development. The reflex evaluates the ability of a mouse to turn themselves from side-position to prone-position (preferred). The scoring, done with visual inspection, is based on the time it takes to return to prone position, and is as follows (with numerical value): 0-2 seconds (6); 2-4 seconds (5); 4-7 seconds (4); 7-10 seconds (3); 10-15 seconds (2); 15-30 seconds (1); and > 30 seconds (0). Testing was performed at P3, P6, P9, and P12.Lr

### Locomotor Movement

Typically developing mice crawl from birth to five days postnatal age, and transition to walking by 10 days postnatally. This sensory-motor assessment places a mouse in an open field for two mins and evaluates movement. The scoring is as follows (with numerical score): fast crawling/walking (3); crawling with symmetric limb movement (2); slow crawling with asymmetric limb movement (1); and no movement (0). Annotations are made for symmetric limb movement (hindlimbs meet front paws during each step with smooth transition between steps) and asymmetric limb movement (inconsistent paw placement and irregular stepping pattern). Testing was performed at P3, P6, P9, and P12.Lr

### Hindlimb Foot Angle

Hindlimb foot angle is an indicator of both sensory-motor and motor development. A picture is taken while a mouse is ambulating in a straight line. Then, the hindlimb foot angle is measured by superimposing a line from the heel to the tip of the middle toe for each foot and identifying the angle made by the dorsal intersection of these two lines. This evaluation identifies abnormal development of the hindlimb as the mouse matures from crawling to walking. In the typical case, the hindlimb angle is reduced in walking compared to crawling. Testing was performed at P6, P9, and P12.Lr

### Hindlimb Strength

The hindlimb suspension test is used to hindlimb strength. Mice are positioned with their hindlimbs at the rim of a 50mL conical tube. Pads are placed at the bottom of the conical tube to cushion a fall, if needed. Testing is performed with two attempts to determine if a mouse can (with numerical score): hold the position for three seconds (4); weakly maintain the position (3); touch the tube with inconsistency (2); clasp hindlimbs to maintain position (1); or fail to hold the position during any of the attempts (0). Secondary measures included latency to fall (number of seconds). Testing was performed at P6, P9, and P12.

### Forelimb Strength

The forelimb suspension test is used to evaluate forelimb and paw strength. Mice are positioned with their forelimbs grasping the rim of a 50mL conical tube. Pads are placed at the bottom of the conical tube to cushion a fall, if needed. Testing is performed with two attempts to determine (with numerical score): normal limb muscle tone (2); muscle weakness (1); or loss of muscle tone or paralysis (0). Secondary measures included latency to fall (number of seconds). Testing was performed at P6, P9, and P12.

### Open Field Test

The open field testing examines basic locomotion levels and anxiety-like behavior. Each mouse is gently placed in a corner of an open field Plexiglas clear chamber (21 cm x 21 cm x 30 cm) and allowed to move freely for 30 mins. Locomotion data is collected using the open field activity monitoring system (AccuScan Instruments, Inc. Columbus, OH), which uses photocell emitters and receptors forming an x-y grid of invisible infrared beams at the base of the chambers. As animals move, the analyzer records the beams break information to observe the animal’s locomotion patterns. Testing was performed from P35 - P40.

### Social Interaction Assay

The socialization chamber, used to evaluate social deficits in mice, is composed of three partitions separated by retractable doorways. A single mouse is placed in the middle chamber and allowed to explore freely. After 10 mins, the mouse is given free access to the other two chambers for another 10 mins. After this exploratory period, an inverted empty wire cup (novel object) is placed in one of the chambers, and an unfamiliar (novel) sex- and age-matched mouse (i.e., males with males and females with females) is placed inside an identical inverted wire cup in the other chamber. To control for the possibility of innate side preferences, the wire cups with and without the novel mouse are randomly placed on the left or right chambers, and the placement is changed between mice. After the acclimation period, mice are video recorded for 10 mins. Total time spent sniffing the novel object, novel mouse, as well as the total time spent in each chamber are recorded and scored. Testing was performed from P40 - P45.

### Rotarod

An accelerating Rotarod apparatus (TSE Instruments) assesses learning, balance, and motor coordination. Rotarod testing involves placing mice on a rotating bar and determining the length of time that they can retain their balance as the rate of rotation is increased. Mice complete one five-minute acclimation session at 15 rotations per minute (RPM) prior to the testing phase. They then undergo two trials (5 mins per trial, 30-minute break between trials) per day for four consecutive days (Max 40 rpm, ∼0.2 rpm increase/second). The latencies to fall from the rotarod are recorded. Testing was performed from P50 - P55.

### Y-Maze

The Y-Maze test evaluates short-term memory and cognition in mice by observing tendencies to explore new areas or stay in a familiar place. The maze consists of three arms (labeled A, B, and C). Mouse movements are tracked over a ten-minute session, and steps are recorded in sets of three to analyze three parameters. Spontaneous Alternation Performance (SAP) measure sequences of three consecutive visits to different arms (ABC, BCA, etc.). Frequency of Alternation Returns (AAR) records non-consecutive same-arm returns (BAB, ACA, etc.). Same Arm Returns (SAR) documents consecutive same-arm returns (BBA, CAA, etc.). No acclimation period was needed. Each mouse underwent one trial. Testing was performed from P116 - P120.

### Elevated Plus Maze

The elevated plus maze is used to quantify anxiety levels. The maze consists of four arms: two closed arms, and two open arms. Anxious mice usually spend more time on the closed arms, and non-anxious mice tend to explore the open arms more. Mice are placed in the center of the maze for 5 mins of testing. The frequency and duration of entries into each arm are recorded. No acclimation period is needed. Each mouse underwent one trial. Testing was performed from P123 - P130.

### MRI

MRI was used to investigate the effects of prenatal viral infection on the adult brain. Adult mice were euthanized with isoflurane. After perfusion with PFA and decapitation, mouse heads were stored in PFA and transferred from Children’s National Hospital to Howard University for imaging. MRI was performed on a preclinical 9.4T 89mm vertical bore Avance400 system (Bruker BioSpin Corp., Billerica, MA). The mouse heads were placed in a glass vial, immersed in perfluorinaded fluid (Galden LS 200, Solvay Specialty Polymers, Alpharetta, GA), and packed with cotton gauze for stability. Whole-brain images were acquired with a 25mm inner diameter RF volume coil. T2-weighted 3D RARE (Rapid Acquisition with Refocused Echoes) fast spin echo anatomical images (TE_eff_ = 36ms, TR = 1500ms, Echo Train Length = 8, FOV = 1.7×1.28×0.72cm, Matrix = 340×256×72, Averages = 10) and 3D spin echo DTI (Diffusion Tensor Imaging) images (TE = 26.5ms, TR=275ms, 15 directions, 1 A0 image, single shell, B = 1000s/mm2, FOV = 1.7×1.4×1.4cm, Matrix = 128×96×96, Averages = 1) were acquired. Imaging was performed in adulthood, with an age range of five to eight months.

### Statistical Analysis

All statistics were completed using GraphPad Prism software. All mice were treated independently, even though litter affected outcome in some cases. All data was first assessed for normality. Normal data was analyzed using one-way ANOVA, two-way ANOVA, two-way ANOVA with mixed effects model, Fisher’s Exact test, individual unpaired t-test, one-tailed student t-test, or simple linear regression. Nonparametric data was analyzed using the Mann-Whitney test or the Kruskal-Wallis test with Dunn’s multiple comparisons. All error bars display the standard error of the mean. Some error bars are not shown because standard error of the mean is smaller than the space covered by the data point. Statistical outliers (more than two standard deviations away from the mean) were omitted occasionally. Figure legends indicate the biggest sample sizes (n) used. We reported P values in the New England Journal of Medicine (NEJM) style. A P value < 0.05 was considered statistically significant; we used (*) when P < 0.05; (**) when P < 0.01; and (***) when P < 0.001. All individual tests are described in supplemental files.

### BioRender

Icons used in figures 1, 4, 5, and 6 and supplemental figures 1 and 2 were available to the authors through an institutional license to Biorender.com.

## MISCELLANEOUS

### Author Contributions

A.R.H. and Y.A.K. conceived and designed the study. A.R.H., C.M.A., and Y.A.K. wrote the manuscript. A.R.H., C.M.A., and E.P. performed experiments and data analyses. S.A.M. completed the sex analysis. C.M.A., A.P.A., A.H.P., N.G.P., J.E.R., A.T.A., and E.C.L. performed tissue sectioning and staining. C.M.A., A.P.A., K.E.H., A.H.P., J.E.R., and A.T.A. completed blinded measurements. S.L. and P.C.W. performed MRI imaging. L.W. completed the open field test, Y-Maze test, and the elevated plus maze test. H.A.G.D. performed statistical analyses. T.F.H., Z.L., and T.A.M. contributed to study design and data interpretation. Y.A.K. supervised the research project.

## Acknowledgments

We would like to thank the faculty and staff at the Centers for Genetic Medicine and Neuroscience Research at Children’s National Research Institute, as well as the Pathology and Neurobehavioral Evaluation Cores at Children’s National Hospital for their support. We would also like to thank Dr. Michael S. Diamond, Dr. Arianna L. Smith, Dr. Lakshmi R. Nair, Ms. Caroline G. Hobson, Ms. Renee A. Jaranson, and Ms. Ashley J. Pocasangre for their contributions.

## REFERENCES

1. Pielnaa, P. et al. Zika virus-spread, epidemiology, genome, transmission cycle, clinical manifestation, associated challenges, vaccine and antiviral drug development. Virology 543, 34–42 (2020).

2. Chiu, C.F. et al. The Mechanism of the Zika Virus Crossing the Placental Barrier and the Blood-Brain Barrier. Front Microbiol 11, 214 (2020).

3. The history of zika virus (Accessed July 10, 2024). (World Health Organization, 2016).

4. Freitas, D.A., et al. Congenital Zika syndrome: A systematic review. PLoS One 15, e0242367 (2020).

5. Woodson, S.E. & Morabito, K.M. Continuing development of vaccines and monoclonal antibodies against Zika virus. NPJ Vaccines 9, 91 (2024).

6. Grant, A. et al. Zika Virus Targets Human STAT2 to Inhibit Type I Interferon Signaling. Cell Host Microbe 19, 882–90 (2016).

7. Kumar, A. et al. Zika virus inhibits type-I interferon production and downstream signaling. EMBO Rep 17, 1766–1775 (2016).

8. Caine, E.A., Jagger, B.W. & Diamond, M.S. Animal Models of Zika Virus Infection during Pregnancy. Viruses 10(2018).

9. Shelton, S.M. et al. Forebrain Neural Precursor Cells Are Differentially Vulnerable to Zika Virus Infection. eNeuro 8(2021).

10. Mesci, P. et al. Blocking Zika virus vertical transmission. Sci Rep 8, 1218 (2018).

11. Julander, J.G. et al. Consequences of in utero exposure to Zika virus in offspring of AG129 mice. Sci Rep 8, 9384 (2018).

12. Li, P. et al. Zika Virus Attenuation by Codon Pair Deoptimization Induces Sterilizing Immunity in Mouse Models. J Virol 92(2018).

13. Cugola, F.R. et al. The Brazilian Zika virus strain causes birth defects in experimental models. Nature 534, 267–71 (2016).

14. Sapparapu, G. et al. Neutralizing human antibodies prevent Zika virus replication and fetal disease in mice. Nature 540, 443–447 (2016).

15. Regla-Nava, J.A. et al. Cross-reactive Dengue virus-specific CD8(+) T cells protect against Zika virus during pregnancy. Nat Commun 9, 3042 (2018).

16. Shi, Y. et al. Vertical Transmission of the Zika Virus Causes Neurological Disorders in Mouse Offspring. Sci Rep 8, 3541 (2018).

17. Miner, J.J. et al. Zika Virus Infection during Pregnancy in Mice Causes Placental Damage and Fetal Demise. Cell 165, 1081–1091 (2016).

18. Yockey, L.J. et al. Vaginal Exposure to Zika Virus during Pregnancy Leads to Fetal Brain Infection. Cell 166, 1247–1256 e4 (2016).

19. Wu, K.Y. et al. Vertical transmission of Zika virus targeting the radial glial cells affects cortex development of offspring mice. Cell Res 26, 645–54 (2016).

20. Shao, Q. et al. The African Zika virus MR-766 is more virulent and causes more severe brain damage than current Asian lineage and dengue virus. Development 144, 4114–4124 (2017).

21. Cui, L. et al. Visual and Motor Deficits in Grown-up Mice with Congenital Zika Virus Infection. EBioMedicine 20, 193–201 (2017).

22. Gorman, M.J. et al. An Immunocompetent Mouse Model of Zika Virus Infection. Cell Host Microbe 23, 672–685 e6 (2018).

23. Hindle, S. et al. Zika virus infection during pregnancy and vertical transmission: case reports and peptide-specific cell-mediated immune responses. Arch Virol 169, 32 (2024).

24. Melo, A.S. et al. Congenital Zika Virus Infection: Beyond Neonatal Microcephaly. JAMA Neurol 73, 1407–1416 (2016).

25. Driggers, R.W. et al. Zika Virus Infection with Prolonged Maternal Viremia and Fetal Brain Abnormalities. N Engl J Med 374, 2142–51 (2016).

26. Besnard, M. et al. Congenital cerebral malformations and dysfunction in fetuses and newborns following the 2013 to 2014 Zika virus epidemic in French Polynesia. Euro Surveill 21(2016).

27. Honein, M.A. et al. Birth Defects Among Fetuses and Infants of US Women With Evidence of Possible Zika Virus Infection During Pregnancy. JAMA 317, 59–68 (2017).

28. Brasil, P. et al. Zika Virus Infection in Pregnant Women in Rio de Janeiro. N Engl J Med 375, 2321–2334 (2016).

29. Carvalho-Sauer, R., Costa, M., Barreto, F.R. & Teixeira, M.G. Congenital Zika Syndrome: Prevalence of low birth weight and associated factors. Bahia, 2015-2017. Int J Infect Dis 82, 44–50 (2019).

30. Mulkey, S.B. et al. Sequential Neuroimaging of the Fetus and Newborn With In Utero Zika Virus Exposure. JAMA Pediatr 173, 52–59 (2019).

31. Alves, L.V., Hazin, A.N. & Alves, J.G.B. Neuroimaging in Children Born With Congenital Zika Syndrome: A Cohort Study. J Child Neurol 36, 1066–1070 (2021).

32. Rose, C.E. et al. Early Growth Parameters as Predictors of Developmental Delay Among Children Conceived During the 2015-2016 Zika Virus Outbreak in Northeastern Brazil. Trop Med Infect Dis 5(2020).

33. Schuler-Faccini, L. et al. Neurodevelopment in Children Exposed to Zika in utero: Clinical and Molecular Aspects. Front Genet 13, 758715 (2022).

34. Mulkey, S.B. et al. Neurodevelopmental Outcomes of Normocephalic Colombian Children with Antenatal Zika Virus Exposure at School Entry. Pathogens 13(2024).

35. Kousa, Y.A. & Hossain, R.A. Causes of Phenotypic Variability and Disabilities after Prenatal Viral Infections. Trop Med Infect Dis 6(2021).

36. Zhao, Z. et al. Zika Virus Infection Leads to Variable Defects in Multiple Neurological Functions and Behaviors in Mice and Children. Adv Sci (Weinh) 7, 1901996 (2020).

37. Elmore, S.A. et al. Histology Atlas of the Developing Mouse Placenta. Toxicol Pathol 50, 60–117 (2022).

38. McEwan, F., Glazier, J.D. & Hager, R. The impact of maternal immune activation on embryonic brain development. Front Neurosci 17, 1146710 (2023).

39. Alippe, Y. et al. Fetal MAVS and type I IFN signaling pathways control ZIKV infection in the placenta and maternal decidua. J Exp Med 221(2024).

40. Soares de Oliveira-Szejnfeld, P., et al. Congenital Brain Abnormalities and Zika Virus: What the Radiologist Can Expect to See Prenatally and Postnatally. Radiology 281, 203–18 (2016).

41. Abrams, R.P.M. et al. Therapeutic candidates for the Zika virus identified by a high-throughput screen for Zika protease inhibitors. Proc Natl Acad Sci U S A 117, 31365–31375 (2020).

42. Neelam, V. et al. Outcomes up to age 36 months after congenital Zika virus infection-U.S. states. Pediatr Res 95, 558–565 (2024).

43. Di Bella, D.J. et al. Molecular logic of cellular diversification in the mouse cerebral cortex. Nature 595, 554–559 (2021).

44. Gabriel, E. et al. Recent Zika Virus Isolates Induce Premature Differentiation of Neural Progenitors in Human Brain Organoids. Cell Stem Cell 20, 397–406 e5 (2017).

45. Moadab, G. et al. Prenatal Zika virus infection has sex-specific effects on infant physical development and mother-infant social interactions. Sci Transl Med 15, eadh0043 (2023).

46. Brien, J.D., Uhrlaub, J.L., Hirsch, A., Wiley, C.A. & Nikolich-Zugich, J. Key role of T cell defects in age-related vulnerability to West Nile virus. J Exp Med 206, 2735–45 (2009).

47. Yang, F., Zheng, Q. & Jin, L. Dynamic Function and Composition Changes of Immune Cells During Normal and Pathological Pregnancy at the Maternal-Fetal Interface. Front Immunol 10, 2317 (2019).

48. Rabelo, K. et al. Zika Induces Human Placental Damage and Inflammation. Front Immunol 11, 2146 (2020).

49. Yao, B. et al. Epigenetic mechanisms in neurogenesis. Nat Rev Neurosci 17, 537–49 (2016).

50. Chen, S. & Shenoy, A. Placental Pathology and the Developing Brain. Semin Pediatr Neurol 42, 100975 (2022).

51. Pinato, L. et al. Sleep findings in Brazilian children with congenital Zika syndrome. Sleep 41(2018).

52. Cristina da Silva Rosa, B., Hernandez Alves Ribeiro Cesar, C.P., Paranhos, L.R., Guedes-Granzotti, R.B. & Lewis, D.R. Speech-language disorders in children with congenital Zika virus syndrome: A systematic review. Int J Pediatr Otorhinolaryngol 138, 110309 (2020).

53. Marques, F.J.P., Tran, L., Kousa, Y.A. & Leyser, M. Long-term developmental outcomes of children with congenital Zika syndrome. Pediatr Res (2024).

54. Carvalho, A. et al. Clinical and neurodevelopmental features in children with cerebral palsy and probable congenital Zika. Brain Dev 41, 587–594 (2019).

55. Ball, E.E. et al. Prenatal Zika virus exposure is associated with lateral geniculate nucleus abnormalities in juvenile rhesus macaques. Neuroreport 34, 786–791 (2023).

56. Nguyen, H.N., Qian, X., Song, H. & Ming, G.L. Neural stem cells attacked by Zika virus. Cell Res 26, 753–4 (2016).

57. Alvarez-Buylla, A. & Lim, D.A. For the long run: maintaining germinal niches in the adult brain. Neuron 41, 683–6 (2004).

58. Merkle, F.T., Tramontin, A.D., Garcia-Verdugo, J.M. & Alvarez-Buylla, A. Radial glia give rise to adult neural stem cells in the subventricular zone. Proc Natl Acad Sci U S A 101, 17528–32 (2004).

59. Gabriel, E., Ramani, A., Altinisik, N. & Gopalakrishnan, J. Human Brain Organoids to Decode Mechanisms of Microcephaly. Front Cell Neurosci 14, 115 (2020).

60. Wheeler, A.C. et al. Developmental Outcomes Among Young Children With Congenital Zika Syndrome in Brazil. JAMA Netw Open 3, e204096 (2020).

61. Marques, F.J.P. et al. Neurodevelopmental outcomes in a cohort of children with congenital Zika syndrome at 12 and 24 months of age. Child Care Health Dev 49, 304–310 (2023).

62. Mulkey, S.B. et al. Neurodevelopmental Abnormalities in Children With In Utero Zika Virus Exposure Without Congenital Zika Syndrome. JAMA Pediatr 174, 269–276 (2020).

63. Lean, S.C., Derricott, H., Jones, R.L. & Heazell, A.E.P. Advanced maternal age and adverse pregnancy outcomes: A systematic review and meta-analysis. PLoS One 12, e0186287 (2017).

64. Zheng, G.X. et al. Massively parallel digital transcriptional profiling of single cells. Nat Commun 8, 14049 (2017).

65. Fleming, S.J. et al. Unsupervised removal of systematic background noise from droplet-based single-cell experiments using CellBender. Nat Methods 20, 1323–1335 (2023).

66. Wolf, F.A., Angerer, P. & Theis, F.J. SCANPY: large-scale single-cell gene expression data analysis. Genome Biol 19, 15 (2018).

67. Sievert, C. Interactive web-based data visualization with R, plotly, and shiny. (2019).

68. Adam Gayoso, J.S. JonathanShor/DoubletDetection: doubletdetection v4.3.0.post1. (2025).

69. Xu, C. et al. Automatic cell-type harmonization and integration across Human Cell Atlas datasets. Cell 186, 5876–5891 e20 (2023).

70. Korsunsky, I. et al. Fast, sensitive and accurate integration of single-cell data with Harmony. Nat Methods 16, 1289–1296 (2019).

