## Supplementary Information for "A humanized mouse model system mimics prenatal Zika infection and reveals premature differentiation of neural stem cells"

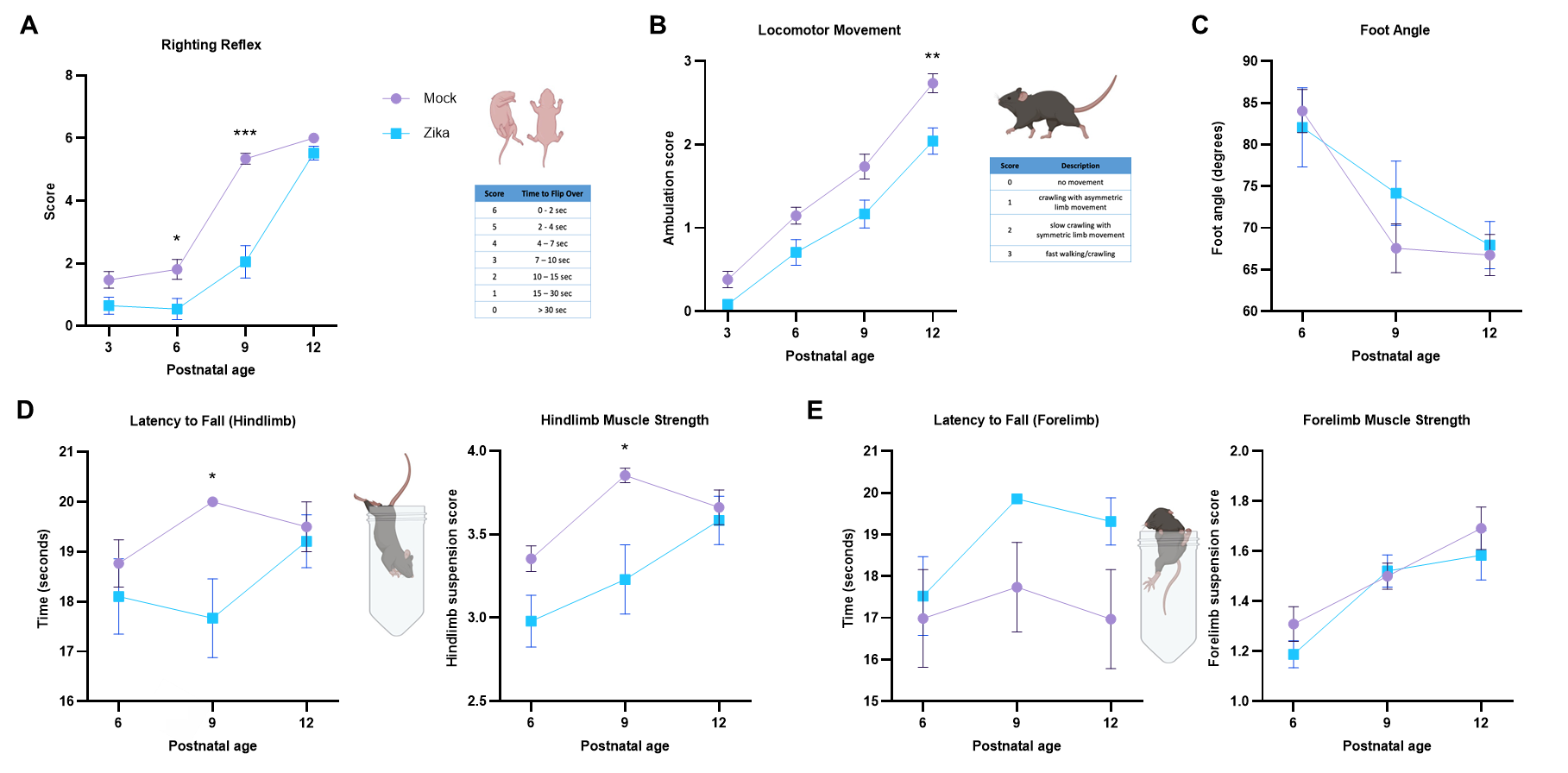

**Supplemental Fig. 1:**

**a-e** Data from Figure 6, only including mice that survived the duration of motor milestone testing (n (litters, embryos) = (2, 12 mock), (4, 12 Zika)). Developmental testing included righting reflex **(a),** locomotor movement **(b),** foot angle while walking **(c),** hindlimb suspension **(d),** and forelimb suspension **(e).** For at least one timepoint, infected mice demonstrated delay in righting reflex, locomotor movement, hindlimb suspension score, and hindlimb strength. There was no significance in foot angle, latency to fall (hindlimb and forelimb), or forelimb suspension score.

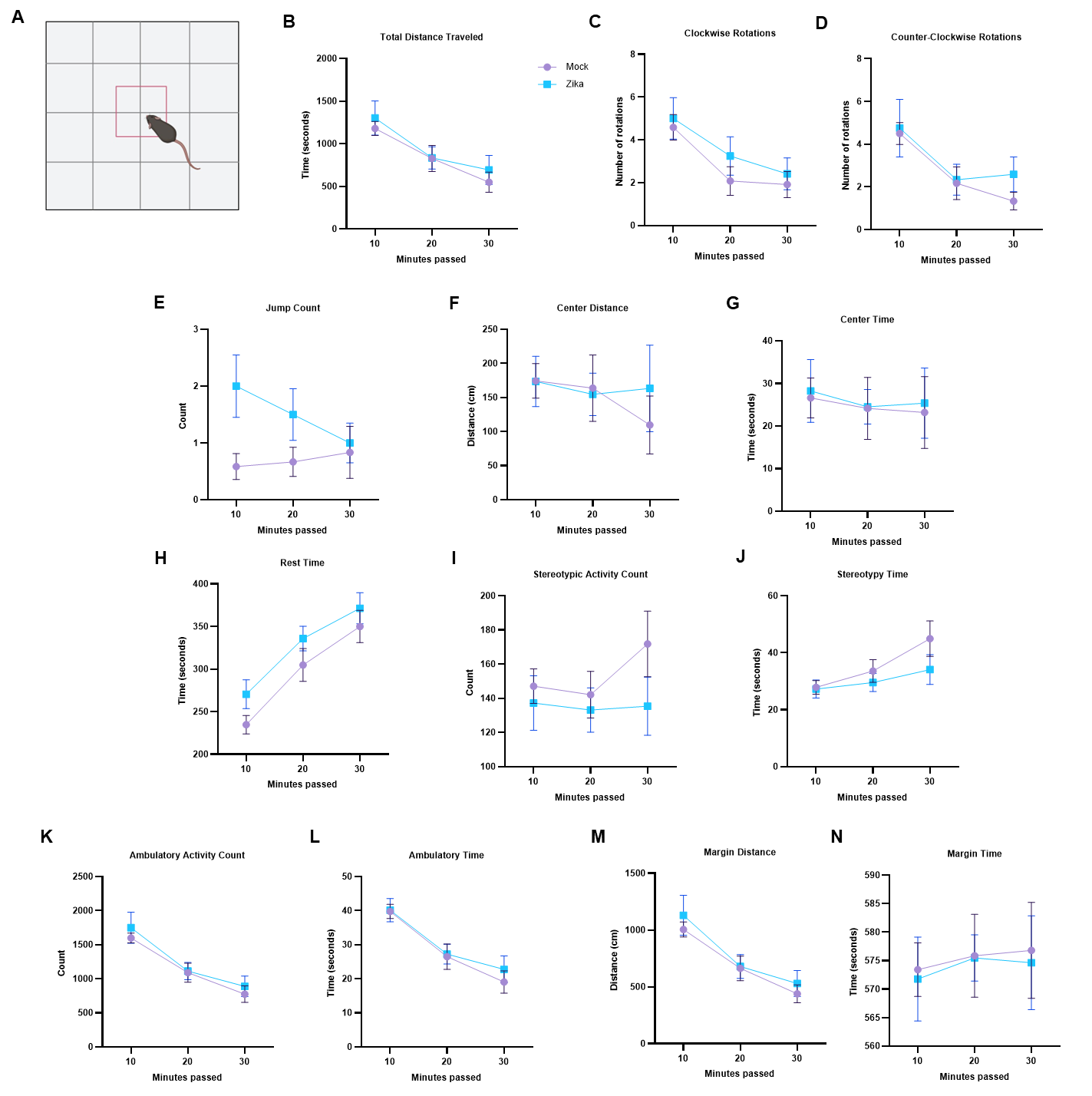

**Supplemental Fig. 2:**

Various ambulatory measurements from mock- or Zika-infected mice in an open field test, which lasted for 30 mins (n (litters, embryos): (2, 12 mock), (2, 12 Zika). There was no significant difference between mock- or Zika-infected mice. We used (*) when P < 0.05; (**) when P < 0.01; and (***) when P < 0.001.

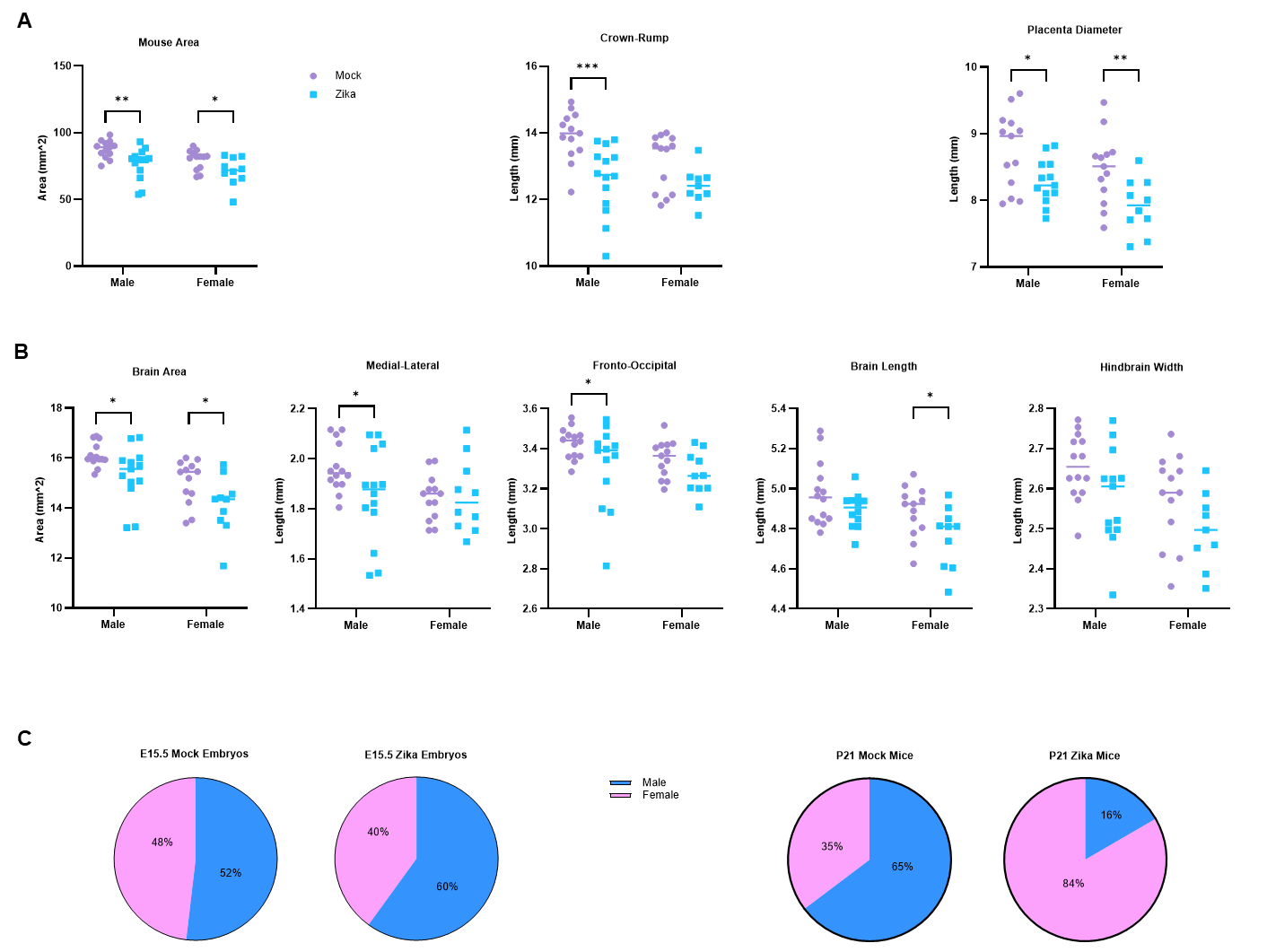

**Supplemental Fig. 3**:

**a-b** Data from Figure 3d-e, separated and evaluated by sex. Sex and infection condition were significant predictors of prenatal gross measurements individually, but corporately, sex and infection condition were not significantly correlated with outcome.

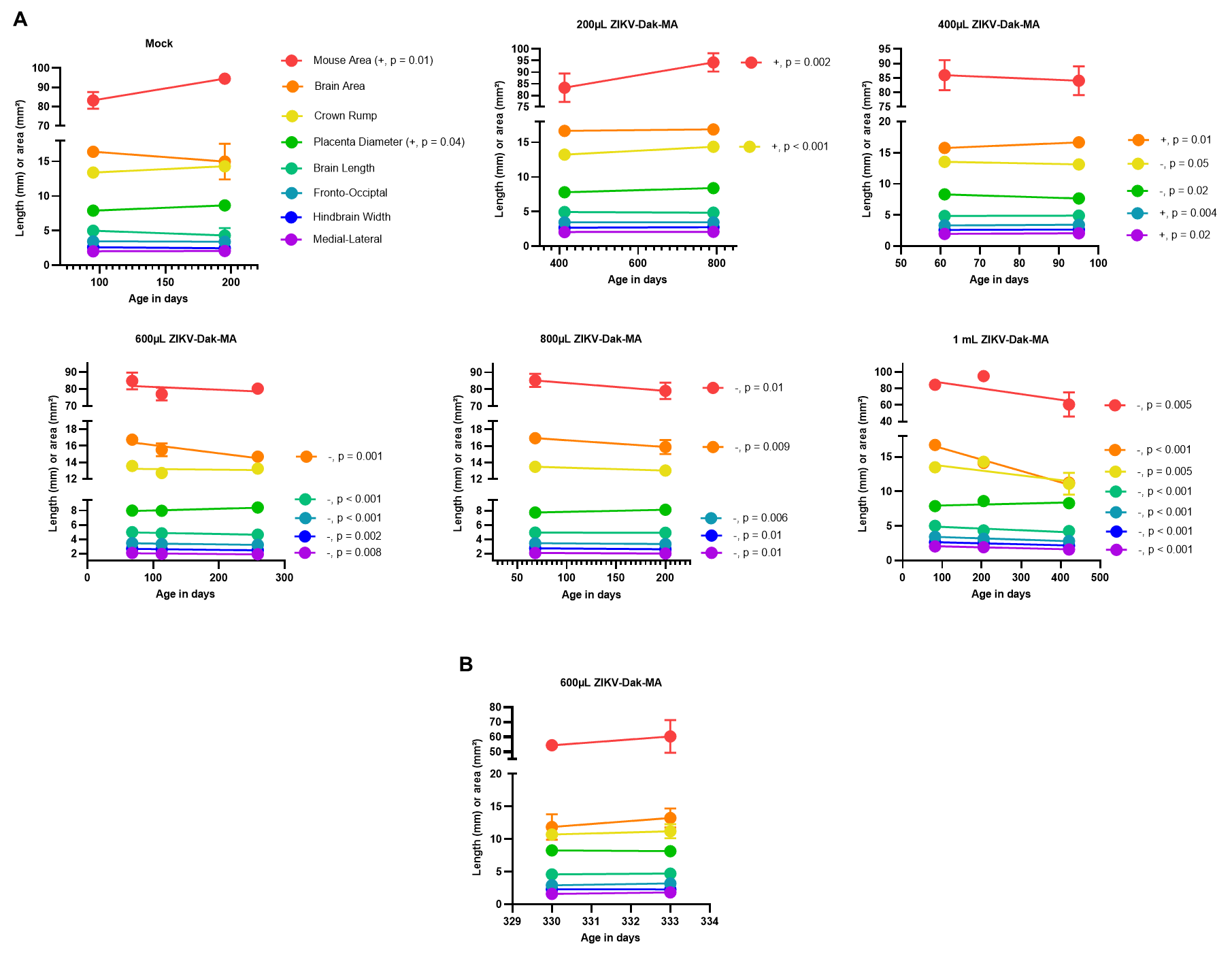

**Supplemental Fig. 4**

**a** E10.5 data from Figure 2a, separated and evaluated by maternal age. Advanced maternal age contributed to adverse prenatal outcomes, particularly at higher viral doses. **b** E9.5 data from Figure 2d-e, separated and evaluated by maternal age. Advanced maternal age contributed to adverse prenatal outcomes, particularly at higher viral doses. We used (*) when P < 0.05; (**) when P < 0.01; and (***) when P < 0.001.

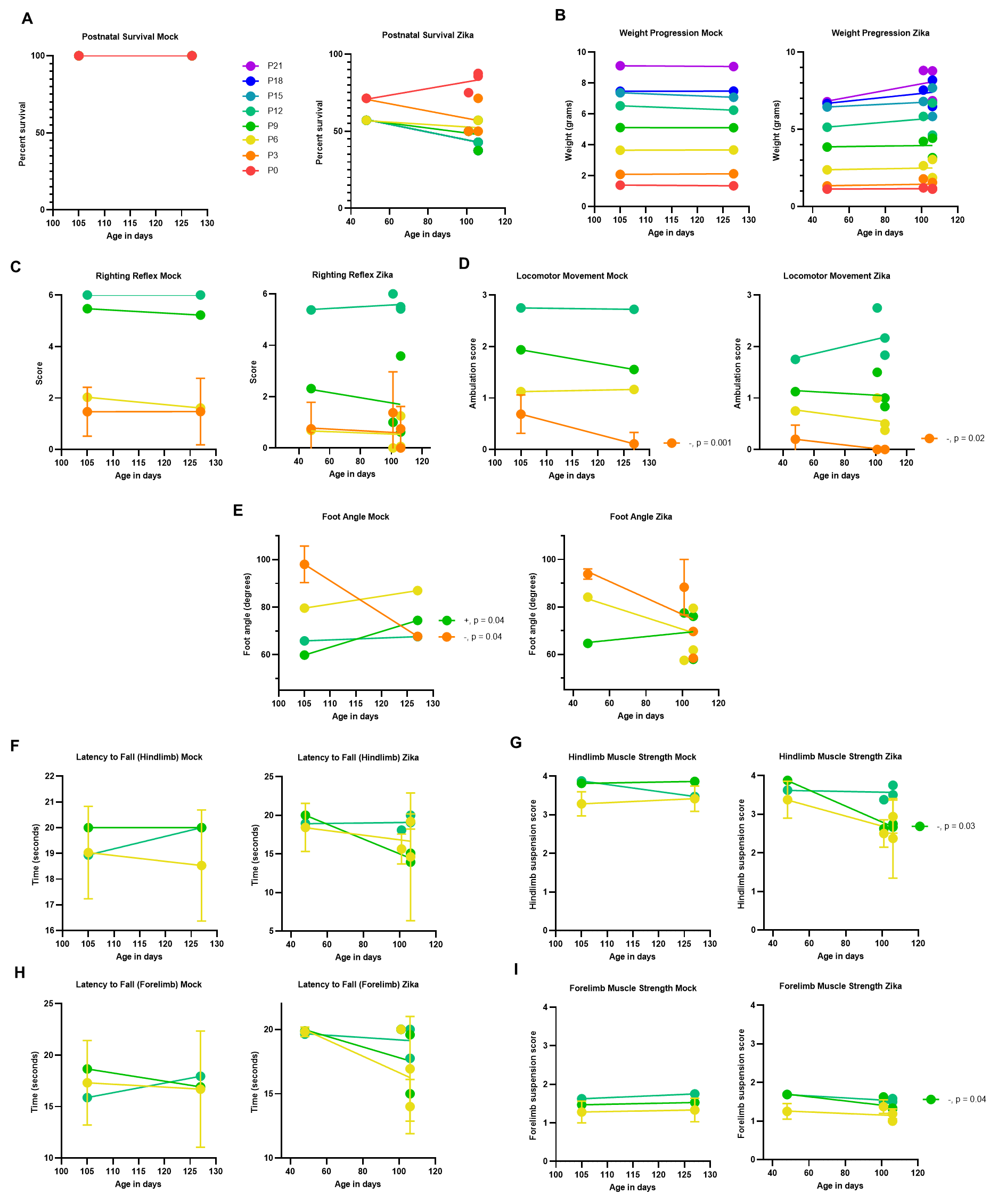

**Supplemental Fig. 5**

Data from Figure 6, separated and evaluated by maternal age (maternal age, # of embryos in the litter) = Mock: (127, 9), (105, 8). Zika: (48, 5), (106, 7), (106, 6) (101, 3). Maternal age was not a modifier for postnatal outcomes.

**Supplemental Statistical Information:**

**Fig. 1b**: E12.5 data were analyzed with the Kruskal-Wallis test, corrected with Dunn’s multiple comparisons tests, and are as follows: n (litters, embryos) = (1, 5 mock), (1, 9 Zika). Zika vs. PC: Placenta: P value < 0.001. Cortex: P < 0.001. Hindbrain: P < 0.001.

**Fig. 1c**: E15.5 data were analyzed with Kruskal-Wallis test and are as follows: Maternal saliva: n (litters, embryos) = (3, 3 mock), (8, 8 Zika). P = 0.02. Placenta: n = (3, 8 mock), (2, 13 Zika). P < 0.001. Fetal blood: n = (2, 4 mock), (3, 5 Zika). P = 0.008. Fetal brain: n = (2, 2 mock), (2, 6 Zika). P = 0.05.

**Fig. 1e**: Data were analyzed with individual unpaired t-tests and are as follows: n (litters, embryos) = (2, 5 mock), (2, 4 Zika). Total area: P = 0.0476. Decidua: P = 0.0317. Junctional zone: P = 0.2038. Labyrinth: P = 0.0588. Chorionic plate: P = 0.0094.

**Fig. 2a**: Data were analyzed with a one-way ANOVA and Dunnett’s test to correct for multiple comparisons and are as follows: DPBS: n (litters, embryos) = (3, 12). Viral titer in PFU: 7.78 x 10^7^, 1.56 x 10^8^, 2.33 x 10^8^, 3.11 x 10^8^, and 3.89 x 10^8^. Virus: n = (2, 15); (2, 15); (3, 18); (3, 18); and (3, 18). ANOVA Summary: P value = 0.03, F = 2.723, DFn, DFd = 5, 82.

**Fig. 2b**: Data were analyzed by a Cochran-Armitage Test and are as follows: DPBS: n (litters, embryos) = (3, 10). Virus: n = (2, 14), (2, 15), (3, 18), (3, 18), and (3, 15). P = 0.058.

**Fig. 2d-e:** Data were analyzed with two-way ANOVAs with mixed effect method, corrected using Šídák's multiple comparisons test, and are as follows: Placenta Diameter: Timepoint of Infection: P = 0.13. F = 2.337. DFn, DFd = 1, 77. IC: P = 0.004. F = 8.768. DFn, DFd = 1, 77. Timepoint of Infection x IC: P = 0.15. F = 2.130. DFn, DFd = 1, 77. Mouse Area: Timepoint of Infection: P = 0.01. F = 6.736. DFn, DFd = 1, 76. IC: P < 0.001. F = 16.04. DFn, DFd = 1, 76. Timepoint of Infection x IC: P = 0.53. F = 0.4053. DFn, DFd = 1, 76. Crown-Rump: Timepoint of Infection: P = 0.01. F = 7.051. DFn, DFd = 1, 75. IC: P < 0.001. F = 18.70. DFn, DFd = 1, 75. Timepoint of Infection x IC: P = 0.21. F = 1.604. DFn, DFd = 1, 75. Brain Area: Timepoint of Infection: P = 0.006. F = 7.917. DFn, DFd = 1, 77. IC: P = 0.06. F = 3.573. DFn, DFd = 1, 77. Timepoint of Infection x IC: P = 0.25. F = 1.349. DFn, DFd = 1, 77. Brain Length: Timepoint of Infection: P = 0.16. F = 2.026. DFn, DFd = 1, 77. IC: P = 0.78. F = 0.08149. DFn, DFd = 1, 77. Timepoint of Infection x IC: P = 0.07. F = 3.441. DFn, DFd = 1, 77. Hindbrain Width: Timepoint of Infection: P = 0.91. F = 0.01159. DFn, DFd = 1, 77. IC: P = 0.52. F = 0.4263. DFn, DFd = 1, 77. Timepoint of Infection x IC: P = 0.02. F = 5.486. DFn, DFd = 1, 77. Medial-Lateral: Timepoint of Infection: P = 0.03. F = 4.714. DFn, DFd = 1, 77. IC: P = 0.01. F = 6.754. DFn, DFd = 1, 77. Timepoint of Infection x IC: P = 0.49. F = 0.4718. DFn, DFd = 1, 77. Fronto-Occipital: Timepoint of Infection: P = 0.001. F = 11.13. DFn, DFd = 1, 77. IC: P < 0.001. F = 12.49. DFn, DFd = 1, 77. Timepoint of Infection x IC: P = 0.98. F = 0.0007772. DFn, DFd = 1, 77.

**Fig. 3d-f**: Data were analyzed with Mann-Whitney tests and are as follows: Forebrain (Region 1): n (litters, embryos) = (2, 6 mock), (2, 9 Zika). Lateral Ventricle: P value = 0.8639. Germinal Zone: P = 0.005. Cortical Plate: P = 0.066. Hemisphere: P = 0.026. Anterior Midbrain (Region 2): n = (2, 5 mock), (2, 7 Zika). Lateral Ventricle: P = 0.530. Germinal Zone: P = 0.01. Cortical Plate: P = 0.048. Midbrain: P = 0.03. Hemisphere: P = 0.03. Posterior Midbrain (Region 3): n = (2, 6 mock), (1, 6 Zika). Lateral Ventricle: P = 0.589. Germinal Zone: P = 0.937. Cortical Plate: P = 0.589. Midbrain: P = 0.818. Hemisphere: P = 0.485.

**Fig. 4**: For all measurements in figure 4, n (litters, embryos) = P0: (2, 17 mock), (4, 21 Zika). P3: (2, 17 mock), (4, 16 Zika). P6: (2, 17 mock), (4, 14 Zika). P9: (2, 17 mock), (4, 13 Zika). P12: (2, 17 mock), (4, 12 Zika).

**Fig. 4b**: Data were analyzed with a log-rank (Mantel-Cox) test and are as follows: Chi square = 9.640, DF = 1, P value = 0.0019.

**Fig. 4c**: Data were analyzed with two-way ANOVAs with mixed effect method with a random coefficient, corrected using Šídák's multiple comparisons test, and are as follows: Fixed Effects: Time: P < 0.0001. F = 824.8. DFn, DFd = 1.555, 43.33. Infection Condition (IC): P < 0.0001. F = 17.47. DFn, DFd = 1, 36. Time x IC: P < 0.0001. F = 6.023. DFn, DFd = 7, 195.

**Fig. 4d-h:** Data were analyzed with two-way ANOVAs with mixed effect method and a random coefficient, corrected using Šídák's multiple comparisons test, and are as follows: Righting Reflex Fixed Effects Results: Time: P < 0.0001. F =130.9. DFn, DFd = 2.532, 97.08. Infection Condition (IC): P < 0.0001. F = 60.27. DFn, DFd = 1, 115. Time x IC: P < 0.0001. F = 11.50. DFn, DFd = 3, 115. Locomotor Movement Fixed Effects Results: Time: P < 0.0001. F = 113.6. DFn, DFd = 2.547, 71.32. IC: P < 0.0001. F = 30.15. DFn, DFd = 1, 31. Time x IC: P = 0.36. F = 1.087. DFn, DFd = 3, 84. Foot Angle Fixed Effects Results: Time: P < 0.0001. F = 12.70. DFn, DFd = 1.939, 132.8. IC: P = 0.47. F = 0.5226. DFn, DFd = 1, 85. Time x IC: P = 0.39. F = 0.9368. DFn, DFd = 2, 137. Latency to Fall Hindlimb Fixed Effects Results: Time: P = 0.1522. F = 2.021. DFn, DFd = 1.623, 44.62. IC: P = 0.0597. F = 3.841. DFn, DFd = 1, 29. Time x IC: P = 0.0703. F = 2.788. DFn, DFd = 2, 55. Hindlimb Muscle Strength Fixed Effects Results: Time: P = 0.0006. F = 9.310. DFn, DFd = 1.773, 48.77. IC: P=0.0034. F = 10.16. DFn, DFd = 1, 29. Time x IC: P = 0.0350. F = 3.567. DFn, DFd = 2, 55. Latency to Fall Forelimb Fixed Effects Results: Time: P = 0.6363. F = 0.4316. DFn, DFd = 1.849, 50.86. IC: P = 0.3660. F = 0.8434. DFn, DFd = 1, 29. Time x IC: P = 0.6733. F = 0.3984. DFn, DFd = 2, 55. Forelimb Muscle Strength Fixed Effects Results: Time: P < 0.0001. F = 19.28. DFn, DFd = 1.869, 51.40. IC: P = 0.2197. F = 1.573. DFn, DFd = 1, 29. Time x IC: P = 0.6481. F = 0.4371. DFn, DFd = 2, 55.

**Fig. 5**: For all measurements in figure 5, n (litters, embryos) = (2, 12 mock), (4, 12 Zika).

**Fig. 5c**: Data were analyzed with individual unpaired t-tests and are as follows: Time spent in each chamber: Novel object chamber: P = 0.5909. T ratio = 0.5455. Center chamber: P = 0.05. T = 2.115. Novel animal chamber: P = 0.28. T = 1.109. Entrances to each chamber: Novel object chamber: P = 0.55. T = 0.6045. Novel animal chamber: P = 0.45. T = 0.7717. Time spent with novelties: Novel object: P = 0.73. T = 0.3445. Novel animal: P = 0.48. T = 0.7212. Entrances to each chamber: Novel object chamber: P = 0.5517. T = 0.6045. Novel animal chamber: P = 0.4485. T = 0.7717. DF for all calculations was 22.00.

**Fig. 5d**: Data were analyzed with two-way ANOVAs with mixed effect method, corrected using Šídák's multiple comparisons test, and are as follows: Rotarod Fixed Effects Results: Time: P = 0.0003. F = 8.458. DFn, DFd = 2.478, 54.52. Infection Condition (IC): P = 0.0263. F = 5.671. DFn, DFd = 1, 22. Time x IC: P = 0.9158. F = 0.1707. DFn, DFd = 3, 66.

**Fig. 5e**: Data were analyzed with individual unpaired t-tests and are as follows: Total steps: P = 0.10. T = 1.741. SAP: P = 0.93. T = 0.08647. AAR: P = 0.09. T = 1.759. SAR: P = 0.08. T = 1.808. DF for all calculations was 22.

**Fig. 5f**: Data were evaluated with individual unpaired t-tests and are as follows: Time spent in each arm: Open: P = 0.35. T = 0.9576. Closed: P = 0.3486. T = 0.9576. Entrances to each arm: Open arm: P = 0.8975. T = 0.1303. Closed arm: P = 0.7161. T = 0.3684. DF for all calculations was 22.

**Fig. 5g**: Data were analyzed with an unpaired t-test and are as follows: P value = 0.05. T value = 2.159, DF = 14.

**Fig. 6d**: Normal data were analyzed with a one-sample Wilcoxon sign rank test or a one-sample t-test and adjusted P values are as follows: Radial glia: Forebrain: P = 0.27. Radial glia: Dorsal Forebrain: P < 0.001. Radial glia: Midbrain: P = 0.059. Neuroblast: Neuronal intermediate progenitor: P < 0.001. Neuroblast: Forebrain GABAergic: P = 0.059. Neuroblast: Thalamus glutamatergic: P = 0.005. Neuroblast: Mixed region: P = 0.028. Neuroblast: Mixed region glutamatergic: P = 0.024. Neuron: Forebrain glutamatergic: P = 0.35. Neuron: Cortical or hippocampal glutamatergic: P < 0.001. Neuron: Forebrain GABAergic: P = 0.018. Neuron: Midbrain GABAergic: P = 0.07. Neuron: Hindbrain GABAergic: P = 0.19. Blood: Erythroid progenitor: P = 0.95. Blood: Erythrocyte: P = 0.021. Blood: Undefined: P = 0.63. Fibroblast: Early fibroblasts: P = 0.43. Immune: Undefined: P = 0.23. Vascular: Undefined: P = 0.52.

**Supplemental Fig. 1**: Data were analyzed using two-way ANOVAs with mixed effect method and corrected with Šídák's multiple comparisons test (Righting Reflex Fixed Effects Results: Time: P value < 0.0001. F =126. DFn, DFd = 2.587, 69.85. Condition: P < 0.0001. F = 51.84. DFn, DFd = 1, 27. Time x Condition: P < 0.0001. F = 9.917. DFn, DFd = 3, 81. Locomotor Movement Fixed Effects Results: Time: P < 0.0001. F = 105.6. DFn, DFd = 2.516, 67.93. Condition: P < 0.0001. F = 23.74. DFn, DFd = 1, 27. Time x Condition: P = 0.4345. F = 0.9210. DFn, DFd = 3, 81. Foot Angle Fixed Effects Results: Time: P < 0.0001. F = 12.70. DFn, DFd = 1.939, 132.8. Condition: P = 0.47. F = 0.5226. DFn, DFd = 1, 85. Time x Condition: P = 0.39. F = 0.9368. DFn, DFd = 2, 137. Latency to Fall Hindlimb Fixed Effects Results: Time: P = 0.2034. F = 1.643. DFn, DFd = 1.970, 53.19. Condition: P = 0.0280. F = 5.392. DFn, DFd = 1, 27. Time x Condition: P = 0.1115. F = 2.285. DFn, DFd = 2, 54. Hindlimb Muscle Strength Fixed Effects Results: Time: P = 0.0007. F = 8.545. DFn, DFd = 1.944, 52.48. Condition: P = 0.0020. F = 11.73. DFn, DFd = 1, 27. Time x Condition: P=0.0773. F = 2.686. DFn, DFd = 1, 54. Latency to Fall Forelimb Fixed Effects Results: Time: P=0.2477. F = 1.434. DFn, DFd = 1.882, 50.82. Condition: P = 0.1168. F = 2.625. DFn, DFd = 1, 27. Time x Condition: P = 0.5629. F = 0.5808. DFn, DFd = 2, 54. Forelimb Muscle Strength Fixed Effects Results: Time: P < 0.0001. F = 19.38. DFn, DFd = 1.802, 48.67. Condition: P = 0.3593. F = 0.8696. DFn, DFd = 1, 27. Time x Condition: P = 0.4737. F = 0.7577. DFn, DFd = 2, 54). Error bars represent standard error of the mean. Some error bars are not shown because standard error of the mean is smaller than the space covered by the data point.

**Supplemental Fig. 2**: Data were analyzed with two-way ANOVAs with mixed effect method, corrected using Šídák's multiple comparisons test, and are as follows: Total Distance Traveled Fixed Effects Results: Time: P value < 0.0001. F = 28.84. DFn, DFd = 1.551, 34.12. Infection Condition (IC): P = 0.6278. F = 0.2417. DFn, DFd = 1, 22. Time x IC: P = 0.6729. F = 0.3998. DFn, DFd = 2, 44. Clockwise Rotation Fixed Effects Results: Time: P value < 0.0001. F = 13.79. DFn, DFd = 1.723, 37.92. IC: P = 0.4403. F = 0.6176. DFn, DFd = 1, 22. Time x IC: P = 0.7423. F = 0.3. DFn, DFd = 2, 44. Counter-Clockwise Rotations Fixed Effects Results: Time: P = 0.0005. F = 12.09. DFn, DFd = 1.393, 30.64. IC: P = 0.5585. F = 0.3530. DFn, DFd = 1, 22. Time x IC: P = 0.6021. F = 0.5132. DFn, DFd = 2, 44. Jump Count Fixed Effects Results: Time: P = 0.5117. F = 0.6324. DFn, DFd = 1.696, 37.3. IC: P = 0.0635. F = 3.818. DFn, DFd = 1, 22. Time x IC: P = 0.1853. F = 1.752. DFn, DFd = 2, 44. Center Distance Fixed Effects Results: Time: P = 0.3993. F = 0.8807. DFn, DFd = 1.553, 34.16. IC: P = 0.7787. F = 0.08093. DFn, DFd = 1, 22. Time x IC: P = 0.4933. F = 0.7181. DFn, DFd = 2, 44. Center Time Fixed Effects Results: Time: P = 0.6967. F = 0.3427. DFn, DFd = 1.862, 40.97. IC: P = 0.8687. F = 0.02799. DFn, DFd = 1, 22. Time x IC: P = 0.9769. F = 0.02339. DFn, DFd = 2, 44. Rest Time Fixed Effects Results: Time: P < 0.0001. F = 44.87. DFn, DFd = 1.908, 41.97. IC: P = 0.1466. F = 2.265. DFn, DFd = 1, 22. Time x IC: P = 0.8206. F = 0.1986. DFn, DFd = 2, 44. Stereotypic Activity Count Fixed Effects Results: Time: P = 0.4680. F = 0.7640. DFn, DFd = 1.933, 42.52. IC: P = 0.2283. F = 1.536. DFn, DFd = 1, 22. Time x IC: P = 0.5092. F = 0.6853. DFn, DFd = 2, 44. Stereotypy Time Fixed Effects Results: Time: P = 0.0128. F = 5.162. DFn, DFd= 1.775, 39.04. IC: P = 0.2217. F = 1.581. DFn, DFd = 1, 22. Time x IC: P = 0.3936. F = 0.9524. DFn, DFd = 2, 44. Ambulatory Activity Count Fixed Effects Results: Time: P < 0.0001. F = 49.47. DFn, DFd = 1.452, 31.94. IC: P = 0.6037. F = 0.2774. DFn, DFd = 1, 22. Time x IC: P = 0.7477. F = 0.2927. DFn, DFd = 2, 44. Ambulatory Time Fixed Effects Results: Time: P < 0.0001. F = 36.71. DFn, DFd = 1.801, 39.62. IC: P = 0.6778. F = 0.1773. DFn, DFd = 1, 22. Time x IC: P = 0.7322. F = 0.3140. DFn, DFd = 2, 44. Margin Distance Fixed Effects Results: Time: P < 0.0001. F = 42.39. DFn, DFd = 1.480, 32.56. IC: P = 0.5944. F = 0.2920. DFn, DFd = 1, 22. Time x IC: P = 0.6942. F = 0.3680. DFn, DFd = 2, 44. Margin Time Fixed Effects Results: Time: P = 0.6965. F = 0.3430. DFn, DFd = 1.862, 40.97. IC: P = 0.8687. F = 0.02798. DFn, DFd = 1, 22. Time x IC: P = 0.9768. F = 0.02345. DFn, DFd = 2, 44.

**Supplemental Fig. 3**: Data were analyzed with two-way ANOVAs and are as follows: n (litters, embryos) = Males: (3, 14 mock), (3, 15 Zika). Females: (3,13 mock), (4, 14 Zika). Mouse Area Source of Variation: Sex: P value = 0.01. F = 6.433. DFn, DFd = 1, 46. Infection Condition (IC): P < 0.001. F = 13.90. DFn, DFd = 1, 46. Interaction: P = 0.67. F = 0.1788. DFn, DFd = 1, 46. Crown-Rump Source of Variation: Sex: P = 0.05. F = 3.954. DFn, DFd = 1, 45. IC: P < 0.001. F = 17.33. DFn, DFd = 1, 45. Interaction: P = 0.22. F = 1.548. DFn, DFd = 1, 45. Placenta Diameter Source of Variation: Sex: P = 0.02. F = 5.594. DFn, DFd = 1, 45. IC: P < 0.001. F = 14.42. DFn, DFd = 1, 45. Interaction: P = 0.79. F = 0.0688. DFn, DFd = 1, 45. Brain Area Source of Variation: Sex: P < 0.001. F = 18.52. DFn, DFd = 1, 45. IC: P = 0.002. F= 10.26. DFn, DFd = 1, 45. Interaction: P = 0.84. F = 0.0430. DFn, DFd = 1, 45. Medial-Lateral Source of Variation: Sex: P = 0.12. F = 2.501. DFn, DFd= 1, 47. IC: P = 0.19. F = 1.784. DFn, DFd = 1, 47. Interaction: P = 0.14. F= 2.311. DFn, DFd = 1, 47. Fronto-Occipital Source of Variation: Sex: P = 0.13. F = 2.344. DFn, DFd = 1, 46. IC: P = 0.02. F = 5.990. DFn, DFd = 1, 46. Interaction: P = 0.58. F = 0.3125. DFn, DFd = 1, 46. Brain Length Source of Variation: Sex: P = 0.006. F = 8.230. DFn, DFd = 1, 45. IC: P = 0.01. F = 7.085. DFn, DFd = 1, 45. Interaction: P = 0.57. F = 0.3352. DFn, DFd = 1, 45. Hindbrain Width Source of Variation: Sex: P = 0.010. F = 7.329. DFn, DFd = 1, 45. IC: P = 0.01. F = 6.744. DFn, DFd = 1, 45. Interaction: P = 0.96. F = 0.0031. DFn, DFd =1, 45. **c** Difference in sex ratios of mock- or Zika-infected litters at E15.5 versus P21. Fisher’s Exact Test demonstrates no statistical difference in sex distribution at E15.5, and a statistically significant difference in distribution between mock- or Zika-infected mice by P21, indicating a survival advantage for the female sex. Data were analyzed with Fisher’s Exact Test are as follows: n (litters, embryos): E15.5: (3, 27 mock), (4, 26 Zika). P21: (2, 17 mock), (2, 12 Zika). E15.5: P = 0.5883. P21: P = 0.0216. We used (*) when P < 0.05; (**) when P < 0.01; and (***) when P < 0.001.

**Supplemental Fig. 4a:** E10.5 data were analyzed with simple linear regressions and are as follows: (maternal age, # of embryos in the litter) = Mock: (95, 6), (195, 3). 200µL ZIKV-Dak-MA: (413, 7), (855, 7). 400µL ZIKV-Dak-MA: (61, 8) (95, 9). 600µL ZIKV-Dak-MA: (68, 7), (113, 7), (259, 4). 800µL ZIKV-Dak-MA: (68, 7), (200, 10). 1mL ZIKV-Dak-MA: (82, 8), (205, 6), (421, 3). Mock: Mouse Area: Y = 0.1126*X + 72.60. P value = 0.01. Brain Area: Y = -0.01425*X + 17.77. P = 0.21. Crown-Rump: Y = 0.009113*X + 12.53. P = 0.08. Placenta Diameter: Y = 0.007636*X + 7.170. P = 0.04. Brain Length: Y = -0.006843*X + 5.651. P = 0.14. Fronto-Occipital: Y = -0.0005833*X + 3.524. P = 0.21. Hindbrain Width: Y = -0.001357*X + 2.735. P = 0.36. Medial-Lateral: Y = 0.0003804*X + 1.993. P = 0.59. 200µL ZIKV-Dak-MA: Mouse Area: Y = 0.02895*X + 71.36. P = 0.002. Brain Area: Y = 0.0005571*X + 16.44. P = 0.52. Crown-Rump: Y = 0.002994*X + 12.00. P < 0.001. Placenta Diameter: Y = 0.001597*X + 7.141. P = 0.05. Brain Length: Y = -0.0002521*X + 5.050. P = 0.19. Fronto-Occipital: Y = -1.738e-005*X + 3.477. P = 0.88. Hindbrain Width: Y = 0.0001854*X + 2.601. P = 0.12. Medial-Lateral: Y = 6.774e-005*X + 2.025. P = 0.51. 400µL ZIKV-Dak-MA: Mouse Area: Y = -0.05654*X + 89.43. P = 0.48. Brain Area: Y = 0.02623*X + 14.18. P = 0.01. Crown-Rump: Y = -0.01217*X + 14.30. P = 0.05. Placenta Diameter: Y = -0.01914*X + 9.489. P = 0.02. Brain Length: Y = 0.001317*X + 4.787. P = 0.21. Fronto-Occipital: Y = 0.003465*X + 3.130. P = 0.004. Hindbrain Width: Y = 0.001654*X + 2.517. P = 0.16. Medial-Lateral: Y = 0.003126*X + 1.793. P = 0.02. 600µL ZIKV-Dak-MA: Mouse Area: Y = -0.01658*X + 82.94. P = 0.34. Brain Area: Y = -0.009911*X + 17.08. P = 0.001. Crown-Rump: Y = -0.0007521*X + 13.27. P = 0.72. Placenta Diameter: Y = 0.002432*X + 7.774. P = 0.10. Brain Length: Y = -0.001675*X + 5.064. P < 0.001. Fronto-Occipital: Y = -0.001200*X + 3.519. P < 0.001. Hindbrain Width: Y = -0.001090*X + 2.728. P = 0.002. Medial-Lateral: Y = -0.0007929*X + 2.092. P = 0.008. 800µL ZIKV-Dak-MA: Mouse Area: Y = -0.04637*X + 88.39. P = 0.01. Brain Area: Y = -0.008086*X + 17.50. P = 0.009. Crown-Rump: Y = -0.003440*X + 13.72. P = 0.09. Placenta Diameter: Y = 0.002894*X + 7.554. P = 0.21. Brain Length: Y = -0.0001603*X + 4.949. P = 0.75. Fronto-Occipital: Y = -0.0009468*X + 3.526. P = 0.006. Hindbrain Width: Y = -0.0009896*X + 2.807. P = 0.01. Medial-Lateral: Y = -0.0006270*X + 2.135. P = 0.01. 1mL ZIKV-Dak-MA: Mouse Area: Y = -0.06872*X + 93.82. P = 0.005. Brain Area: Y = -0.01642*X + 17.90. P < 0.001. Crown-Rump: Y = -0.006905*X + 14.39. P = 0.005. Placenta Diameter: Y = 0.001336*X + 7.838. P = 0.21. Brain Length: Y = -0.002406*X + 5.104. P < 0.001. Fronto-Occipital: Y = -0.001865*X + 3.579. P < 0.001. Hindbrain Width: Y = -0.001397*X + 2.767. P < 0.001. Medial-Lateral: Y = -0.001311*X + 2.172. P < 0.001.

**Supplemental Fig. 4b**: E9.5 data were analyzed with simple linear regressions and are as follows: (maternal age, # of embryos in the litter) = (330, 2), (333, 4). Mouse Area: Y = 1.986*X - 601.0. P = 0.52. Brain Area: Y = 0.4603*X - 140.0. P = 0.42. Crown-Rump: Y = 0.1631*X - 43.10. P = 0.60. Placenta Diameter: Y = -0.03583*X + 20.10. P = 0.83. Brain Length: Y = 0.04250*X - 9.422. P = 0.74. Fronto-Occipital: Y = 0.09383*X - 28.03. P = 0.16. Hindbrain Width: Y = -0.007167*X + 4.690. P = 0.94. Medial-Lateral: Y = 0.07767*X - 24.02. P = 0.06.

**Supplemental Fig. 5a**: Data were analyzed with simple linear regressions and are as follows: Postnatal Survival Mock: For all timepoints: Y = 0.000*X + 100. P value > 0.999. Postnatal Survival Zika: P0: Y = 0.2159*X + 60.41. P = 0.23. P3: Y = -0.2366*X + 82.05. P = 0.46. P6: Y = -0.07850*X + 60.63. P = 0.46. P9: Y = -0.1606*X + 64.92. P = 0.52. P12: Y = -0.2540*X + 69.79. P = 0.16. P15: Y = -0.2540*X + 69.79. P = 0.16. P18: Y = -0.2540*X + 69.79. P = 0.16. P21: Y = -0.2540*X + 69.79. P = 0.16.

**Supplemental Fig. 5b**: Data were analyzed with simple linear regressions and are as follows: Weight Progression Mock: P0: Y = -0.002071*X + 1.607. P = 0.28. P3: Y = 0.001585*X + 1.920. P = 0.56. P6: Y = 0.0008523*X + 3.562. P = 0.91. P9: Y = -0.0001136*X + 5.124. P > 0.99. P12: Y = -0.01271*X + 7.861. P = 0.32. P15: Y = -0.01328*X + 8.765. P 0.34. P18: Y = 0.0004230*X + 7.422. P = 0.98. P21: Y = -0.002203*X + 9.348. P = 0.94. Weight Progression Zika: P0: Y = 0.0004398*X + 1.114. P = 0.65. P3: Y = 0.001836*X + 1.262. P = 0.53. P6: Y = 0.001991*X + 2.291. P = 0.77. P9: Y = 0.001562*X + 3.794. P = 0.88. P12: Y = 0.009943*X + 4.670. P = 0.48. P15: Y = 0.005664*X + 6.178. P = 0.70. P18: Y = 0.01230*X + 6.096. P = 0.38. P21: Y = 0.02157*X + 5.793. P = 0.17.

**Supplemental Fig. 5c**: Data were analyzed with simple linear regressions and are as follows: Righting Reflex Control: P3: Y = 0.0001578*X + 1.452. P > 0.99. P6: Y = -0.01910*X + 4.036. P = 0.53. P9: Y = -0.01121*X + 6.645. P = 0.49. P12: Y = 0.000*X + 6.000. P > 0.999. Righting Reflex Zika: P3: Y = -0.003208*X + 0.9305. P = 0.72. P6: Y = -0.002276*X + 0.7733. P = 0.85. P9: Y = -0.01005*X + 2.762. P = 0.62. P12: Y = 0.003367*X + 5.232. P = 0.70.

**Supplemental Fig. 5d**: Data were analyzed with simple linear regressions and are as follows: Locomotor Movement Control: P3: Y = -0.02620*X + 3.438. P = 0.001. P6: Y = 0.001894*X + 0.9261. P = 0.85. P9: Y = -0.01736*X + 3.760. P = 0.21. P12: Y = -0.001263*X + 2.883. P = 0.91. Locomotor Movement Zika: P3: Y = -0.003490*X + 0.3670. P = 0.02. P6: Y = -0.003976*X + 0.9599. P = 0.52. P9: Y = -0.001709*X + 1.226. P = 0.81. P12: Y = 0.007024*X + 1.439. P = 0.25.

**Supplemental Fig. 5e**: Data were analyzed with simple linear regressions and are as follows: Foot Angle Control: P3: Y = -1.377*X + 242.5. P = 0.04. P6: Y = 0.3350*X + 44.43. P = 0.27. P9: Y = 0.6641*X - 9.893. P = 0.04. P12: Y = 0.08038*X + 57.37. P = 0.69. Foot Angle Zika: P6: Y = -0.3464*X + 111.5. P = 0.17. P9: Y = -0.2459*X + 95.20. P = 0.20. P12: Y = 0.07766*X + 61.28. P = 0.56.

**Supplemental Fig. 5f**: Data were analyzed with simple linear regressions and are as follows: Latency to Fall (Hindlimb) Control: P6: Y = -0.02289*X + 21.43. P = 0.61. P9: Y = 0.000*X + 20.00. P > 0.999. P12: Y = 0.04830*X + 13.87. P = 0.30. Latency to Fall (Hindlimb) Zika: P6: Y = -0.03013*X + 19.83. P = 0.56. P9: Y = -0.09606*X + 24.61. P = 0.10. P12: Y = 0.003002*X + 18.76. P = 0.89.

**Supplemental Fig. 5g**: Data were analyzed with simple linear regressions and are as follows: Hindlimb Muscle Strength Control: P6: Y = 0.006155*X + 2.635. P = 0.40. P9: Y = 0.002210*X + 3.580. P = 0.63. P12: Y = -0.01831*X + 5.797. P = 0.05. Hindlimb Muscle Strength Zika: P6: Y = -0.01297*X + 3.990. P = 0.08. P9: Y = -0.02059*X + 4.857. P = 0.03. P12: Y = -0.0008792*X + 3.659. P = 0.88.

**Supplemental Fig. 5h**: Data were analyzed with simple linear regressions and are as follows: Latency to Fall (Forelimb) Control: P6: Y = -0.02809*X + 20.26. P = 0.80. P9: Y = -0.07907*X + 26.96. P = 0.44. P12: Y = 0.09407*X + 5.998. P = 0.40. Latency to Fall (Forelimb) Zika: P6: Y = -0.06394*X + 23.03. P = 0.06. P9: Y = -0.04176*X + 21.98. P = 0.50. P12: Y = -0.009202*X + 20.10. P = 0.68.

**Supplemental Fig. 5i**: Data were analyzed with simple linear regressions and are as follows: Forelimb Muscle Strength Control: P6: Y = 0.002367*X + 1.033. P = 0.72. P9: Y = 0.002683*X + 1.187. P = 0.59. P12: Y = 0.005682*X + 1.028. P = 0.48. Forelimb Muscle Strength Zika: P6: Y = -0.001988*X + 1.355. P = 0.31. P9: Y = -0.005489*X + 1.960. P = 0.04. P12: Y = -0.002705*X + 1.816. P = 0.49.
